# Host preferences of coexisting Perkinsea parasitoids during coastal dinoflagellate blooms

**DOI:** 10.1101/2021.02.17.431570

**Authors:** Albert Reñé, Natàlia Timoneda, Nagore Sampedro, Elisabet Alacid, Rachele Gallisai, Jordina Gordi, Alan Denis Fernández-Valero, Massimo Ciro Pernice, Eva Flo, Esther Garcés

**Affiliations:** Departament de Biologia Marina i Oceanografia. Institut de Ciències del Mar (CSIC). Pg. Marítim de la Barceloneta, 37-49. 08003 Barcelona, Catalonia (Spain); Living Systems Institute, School of Biosciences, College of Life and Environmental Sciences, University of Exeter. Exeter EX4 4QD (United Kingdom)

**Keywords:** blooms, diversity, interactions, metabarcoding, parasitism, protists

## Abstract

Parasites in aquatic systems are highly diverse and ubiquitous. In marine environments, parasite-host interactions contribute substantially to shaping microbial communities, but their nature and complexity remain poorly understood. In this study, we examined the relationship between Perkinsea parasitoids and bloom-forming dinoflagellate species. Our aim was to determine whether parasite-host species interactions are specific and whether the diversity and distribution of parasitoids are shaped by their dinoflagellate hosts. Several locations along the Catalan coast (NW Mediterranean Sea) were sampled during the blooms of five dinoflagellate species and the diversity of Perkinsea was determined by combining cultivation-based methods with metabarcoding of the V4 region of 18S rDNA. Most known species of Parviluciferaceae, and others not yet described, were detected, some of them coexisting in the same coastal location, and with a wide distribution. The specific parasite-host interactions determined for each of the studied blooms demonstrated the host preferences exhibited by parasitoids in nature. The dominance of a species within the parasitoid community is driven by the presence and abundances of its preferred host(s).The absence of parasitoid species, often associated with a low abundance of their preferred hosts, suggested that high infection rates are reached only under conditions that favour parasitoid propagation, especially dinoflagellate blooms.

## INTRODUCTION

The complexity of natural microbial communities is the product of the diverse interactions of a large number of coexisting species. In marine environments, parasitism has evolved independently in multiple different lineages (Poulin & Randhawa, 2015) and is believed to be one of the main drivers modulating species interactions (de Vargas et al., 2015). However, elucidating the nature and specificity of parasite-host interactions in marine protist communities remains challenging. Phytoplankton play a key role in marine ecosystems as primary producers. Infection by a wide variety of parasitic groups (Skovgaard, 2014), particularly zoosporic parasites has been described as a mortality mechanism of phytoplankton. Zoosporic parasites, representing some distinct lineages, release numerous flagellated zoospores into the environment that actively infect hosts (Jephcott et al., 2016). Because these parasites must kill their hosts to reproduce, they are more accurately referred to as parasitoids. Known zoosporic parasite groups that infect dinoflagellates include chytrids (Frenken et al., 2017; Gleason et al., 2015), Amoebophryidae (Chambouvet et al. 2008; Guillou et al., 2008) and Perkinsea (Jephcott et al., 2016).

The first studies of Perkinsea were conducted in marine systems and focused on *Perkinsus* species, which infect molluscs (Mackin et al. 1950), followed much later by studies of *Parvilucifera*, a parasitoid of dinoflagellates (Norén et al. 1999). In addition to species related to those genera, only a few other species infecting tadpoles (Chambouvet et al., 2015) and marine species of fish (Freeman et al., 2017) have been reported to date. The molecular diversity of Perkinsea has also been explored, but most of those studies focused on freshwater environments, where the parasitoids were found to be relevant in the community and exhibited an unexpected diversity (Bråte et al., 2010; Lefèvre et al., 2008; Lepère et al., 2008; Lepère et al., 2010; Mangot et al., 2011). The use of a molecular TSA-FISH probe in several freshwater environments revealed the presence of Perkinsea in most of the studied lakes and in association with *Sphaerocystis* (Chlorophyceae) (Jobard et al., 2020). However, a study of the presence and interactions of Perkinsea in the South Atlantic Ocean using the same probe failed to detect these parasitoids in the water column (Lepère et al., 2016). Molecular studies of Perkinsea conducted in marine coastal waters revealed most detections in sediment samples (Chambouvet et al. 2014).

Nonetheless, the diversity of Perkinsea remains largely unknown and has thus far mostly been represented by environmental sequences (Chambouvet et al., 2014), although the morphological and phylogenetic features of ten species of parasitoids that infect dinoflagellates have been identified. Those species belong to the genera *Parvilucifera* (Alacid et al., 2020; Figueroa et al., 2008; Jeon et al., 2020; Lepelletier et al., 2014; Norén et al., 1999; Reñé et al., 2017a), *Snorkelia* (Leander & Hoppenrath, 2008; Reñé et al., 2017b), *Dinovorax* (Reñé et al., 2017b) and *Tuberlatum* (Jeon & Park, 2019), all of which are members of the family Parviluciferaceae. Those organisms were described after discrete observations during dinoflagellate blooms from worldwide locations and the successful establishment of parasitoid-host co-cultures.

The presence and infection dynamics of the studied parasitoids were shown to be linked to their hosts. By definition, parasitoids completely depend on their host for survival and reproduction (Waage & Hassell 1982), such that the size of the host population determines their transmission and propagation. Microalgal blooms are characterized by a very simplified phytoplankton community in terms of their diversity, with only one or a few abundant species. Bloom events allow parasitoids to propagate and can be used to examine parasite-host interactions. Indeed, parasitoids are an integral component of the high-density blooms of dinoflagellate species (Park et al., 2013), consistent with the ability of host proliferation to support parasitic epidemics (Alacid et al., 2017). Likewise, Blanquart et al. (2016) used qPCR to demonstrate a correlation between the presence of parasitoids *Parvilucifera infectans* and *P. rostrata* and blooms of the dinoflagellate *Alexandrium minutum*. One of the most important characteristics of parasitoids is their host specificity, which determines their ecology and evolutionary dynamics (Schmid-Hempel, 2011). Host specificity can be measured by host range (number of infected host species), taking into account the prevalence (intensity) and frequency of infections (Rohde, 2002). In turn, host range can be assessed by direct observations in the wild, but these are limited by the need for intensive sampling efforts and the difficulties in parasite-host identification. The use of genetic tools, such as metabarcoding, partially overcomes these limitations, by providing an overview of parasite-host co-occurrences, but the results must then be validated. Host range can also be determined experimentally, which avoids ecological and sampling constraints but only establishes the potential for parasite-host interactions in the lab and again requires validation with observations from nature (Schmid-Hempel, 2011).

Despite studies suggesting the wide host-range of Perkinsea infecting marine phytoplankton (Garcés et al., 2013; Rodríguez & Figueroa, 2020), little is known about its host preferences (Alacid et al., 2016). A complete understanding of the host interactions of Perkinsea requires sampling efforts that capture the infection strategy and life-cycle of these organisms. In this study, we took advantage of recurrent high-biomass blooms of dinoflagellates occurring locally and transiently in Catalan coastal waters to study the interaction of Perkinsea species with their hosts. The structure of the Perkinsea community was explored by sampling both the water column and sediments during high-biomass blooms of target dinoflagellate species, thus ensuring coverage of all possible habitats of parasitoids. As a control, locations where the dinoflagellates are known to recurrently form blooms were also sampled during no-bloom periods. To evaluate parasite-host interactions, we determined whether bloom-forming dinoflagellate species can be infected by different Perkinsea species. Our results, obtained from culture-based methods and environmental metabarcoding, provide insights into parasitoid diversity and distribution and how both are influenced by the host preferences of the parasitoids.

## MATERIALS AND METHODS

### Sampling

The Catalan coast is located in the NW Mediterranean Sea (NE Spain) and extends in a NE– SW orientation from southern France to the southern point of the Ebre Delta (Fig. 1). The sampling stations in this study corresponded to beaches, semi-enclosed waters, such as harbours, and a coastal lagoon situated in a wetland area (Table 1). Samplings were conducted during 2018 and 2019, when blooms of the target dinoflagellates species *Alexandrium minutum*, *A. taylorii*, *Ostreopsis* sp., *Gymnodinium litoralis* and *Kryptoperidinium foliaceum* were detected. Blooms were defined as the presence of the target dinoflagellates at high abundances (> 10^4^ cells L^−1^) in the sampling location (Table 1). For comparison, other locations where the target species are known to recurrently form blooms were sampled during a no-bloom period.

**Table 1.** Characteristics of the sampled locations and sampling date. Each sample was classified based on the main bloom-forming dinoflagellate present in the sample. Dinoflagellate abundances (cells L^−1^) and the contribution of that species (main species, in %) to the dinoflagellate community is presented (from microscopy counts). Asterisks indicate sampling sites where only sediments were processed.

| Code | Location | Type | Ecosystem | Date | Main blooming dinoflagellate | Main species (%) | Abundance (cells L<sup>-1</sup>) |
| --- | --- | --- | --- | --- | --- | --- | --- |
| 1 | El Masnou | Harbour | Marina, the most sensitive harbour by sand accumulation at the Maresme coast | 2-mar-18 | *Alexandrium minutum* | 96 | 3.5 x 10<sup>6</sup> |
| 1b | El Masnou | Harbour |  | 7-mar-18 | *A. minutum* | 94 | 2.2 x 10<sup>6</sup> |
| 2 | Barcelona | Harbour | One of Europe's major ports in the Mediterranean; Catalonia's largest port | 28-mar-19 | *A. minutum* | 90 | 1.3 x 10<sup>5</sup> |
| 3 | Cambrils | Harbour | Small harbour | 15-may-19 | *A. minutum* | 92 | 1.4 x 10<sup>5</sup> |
| 4 | Estartit | Harbour | Small harbour | 5-jun-19 | *A. minutum* | 14 | 9.5 x 10<sup>4</sup> |
| 5 | L'Ametlla de Mar | Harbour | Small harbour | 27-jun-19 | *A. minutum* | 74 | 4.9 x 10<sup>4</sup> |
| 6 | Sant Pol | Beach | Long sand crescent beach, opens up onto a natural bay, about 1 km from end to end, with a wide bay | 25-jul-18 | *Alexandrium taylorii* | 92 | 3.0 x 10<sup>6</sup> |
| 7 | L'Arenal | Beach | Located in Fangar Bay, fine golden sand, typical of the coast of the Ebro Delta | 18-jul-19 | *A. taylorii* | 68 | 4.0 x 10<sup>4</sup> |
| 8 | Aiguafreda | Beach | Semi-urban beach, cobble fragments, calm waters | 4-jul-19 | *A. taylorii* | 64 | 4.7 x 10<sup>6</sup> |
| 9 | Sa Riera * | Beach | Large and sandy beach, north-facing but protected, sand is coarse to grainy | 4-jul-19 | *Gymnodinium litoralis* | 91 | 3.9 x 10<sup>6</sup> |
| 10 | Pals | Beach | Longest beaches on the Costa Brava 3.5 km in total | 4-jul-19 | *G. litoralis* | 96 | 7.4 x 10<sup>6</sup> |
| 11 | Llavaneres | Beach | Rocky beach, urban area | 25-jul-18 | *Ostreopsis* sp. | 100 | 1.9 x 10<sup>4</sup> |
| 12 | Sitges | Beach | Sandy beach, coastal stretch with an ENE-WSW direction; the nearshore sea-bottom of the bay has a regular and low gradient slope, urban area | 7 Aug 2018 | *Ostreopsis* sp. | 92 | $6.0 \times 10^5$ |
| 13 | Cubelles * | Beach | Rocky shore, clear, tranquil waters, detached breakwaters, artificial structures | 21 Aug 2019 | *Ostreopsis* sp. | | $2.5 \times 10^3$ |
| 13b | Cubelles | Beach | | 27 Aug 2019 | *Ostreopsis* sp. | 93 | $5.4 \times 10^4$ |
| 14 | Fra Ramón | Coastal lagoon | Small brackish coastal lagoon located in La Pletera, a salt marsh in the coastal area on the north side of the Ter River mouth, immediately behind the sand dune ridge | 18-jun-19 | *Kryptoperidinium foliaceum* | 41 | $3.9 \times 10^4$ |
| 15 | Vilanova i la Geltrú | Harbour | Small harbour, coastal breakwater | 5-jun-18 | No bloom of target host | 63% *Prorocentrum triestinum* | $2.2 \times 10^6$ |
| 16 | Muga | Beach | Sandy beach, mouth of the La Muga River | 5-jun-19 | No bloom of target host | 45% *P. micans* | total<br>dinoflagellates <<br>$10^4$ |
| 7b | L'Arenal | Beach | Located in Fangar Bay, fine golden sand, typical of the coast of the Ebro Delta | 27-jun-19 | No bloom of target host | 57% *P. triestinum* | total<br>dinoflagellates < 5<br>$\times 10^4$ |

**Figure 1.**
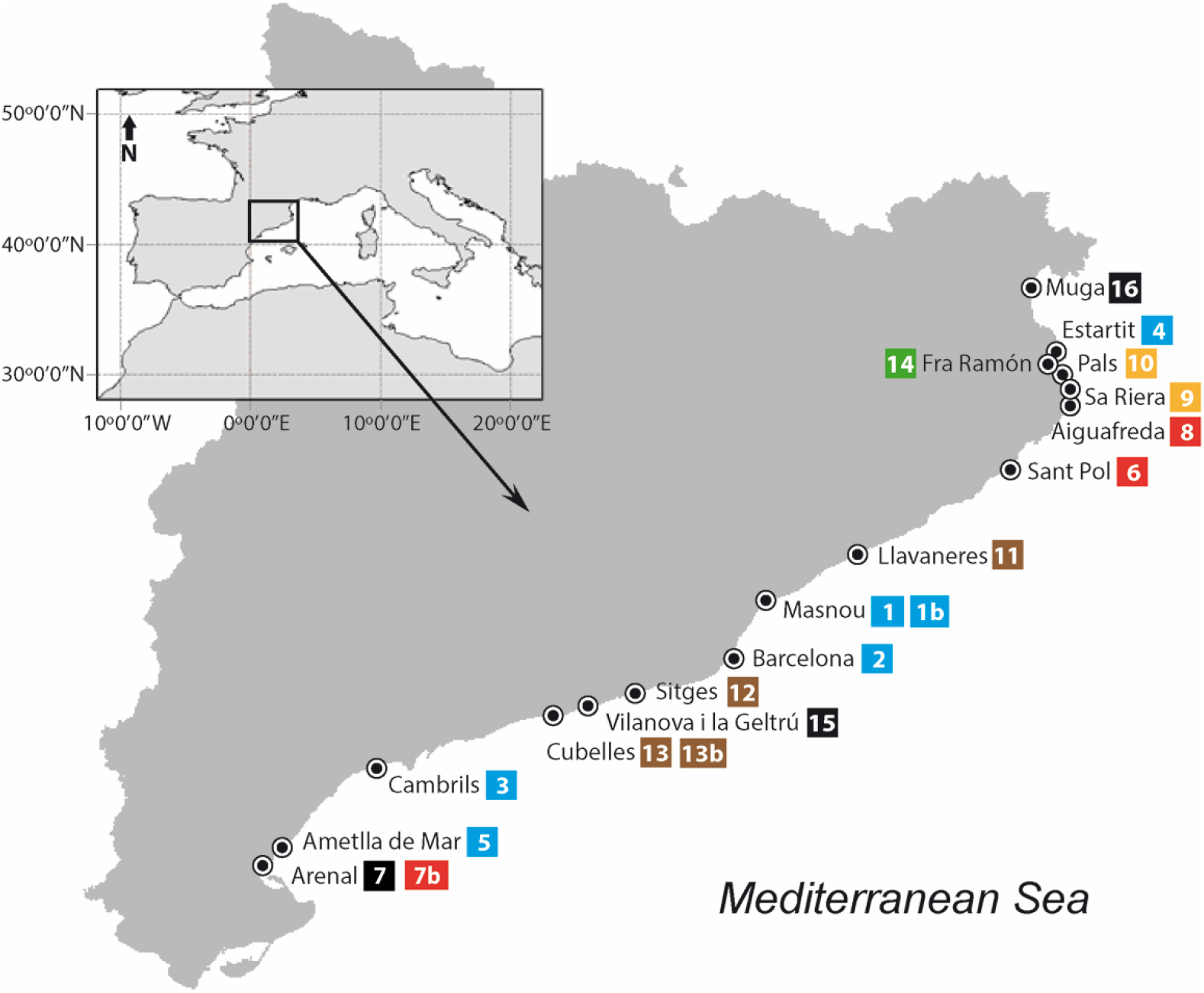
Map of the surveyed areas covering the Catalan coast. A sampling number is assigned to each location. The colours indicate the bloom-forming dinoflagellate species present in the sample: *Alexandrium minutum* (blue), *A. taylorii* (red), *Ostreopsis* sp. (brown), *Gymnodinium litoralis* (yellow), *Kryptoperidinium foliaceum* (green), and no-bloom of target host (black).

Perkinsea parasitoids are present in the water column and sediments. Thus, seawater (~10 L) integrating the whole water column was collected both from harbours (2.5 to 4 m), using a tube connected to an electric pump, and from beaches (~1.5 m depth), where sampling was conducted manually. Seawater was pre-filtered through a 200-μm mesh to remove metazoans and larger particles, after which a 150-mL sample was fixed with Lugol’s iodine for phytoplankton determination. Superficial sediments were obtained snorkelling in waters near the beaches, or by using a pole connected to a sediment corer in harbours. From the sediments, ~30 g were placed in each of two tubes (50 mL) using a spoon. In the laboratory, a portion of the collected seawater was immediately filtered in two replicates by serial filtration through 10-μm and 0.8-μm polycarbonate filters (47 mm) using a peristaltic pump until the filters were saturated (100 mL to 2 L, depending on the total biomass). The filters and sediments were stored frozen at −80°C until processed.

### Microscopic identification and enumeration of the dinoflagellate community

Aliquots of phytoplankton samples (50 mL) previously fixed with Lugol were settled for 24 h in a counting chamber following the Utermöhl method (Utermöhl, 1958). For dinoflagellate enumeration, the appropriate area of the chamber, determined depending on the cell density of each species, was scanned at 200–400× magnification using a Leica-Leitz DMII inverted microscope and at least 30 cells were counted for each species. When needed, the thecal plates of the dinoflagellates were stained with the fluorescent dye Calcofluor white M2R at a final concentration of 10–20 mg mL^−1^ (Fritz & Triemer, 1985) and examined under epifluorescence (lamp 50W) using a Leica DM IRB (Leica Microsystemps GMbH, Wetzlar, Germany) inverted microscope. Sedgewick-rafter counting chambers were used to count cells in the cases of high abundances of a species.

### Detection of infections and the establishment of parasitoid cultures

Crude seawater (5-10 mL) from each location was transferred to 6-well plates and incubated in culture chambers at 21°C with a 10:14 light:dark cycle to allow parasitoids present in the samples to proliferate in their dinoflagellate hosts. For samples containing low dinoflagellate abundances, 1L of seawater was concentrated using a 10-μm net-mesh to increase dinoflagellate and parasitoid abundances and then transferred to the well plates as previously described. Incubations were checked daily especially during the first few days, a period considered crucial for parasitoid survival, by inverted light microscopy in order to detect infections of the dinoflagellates. In samples in which dinoflagellates were not highly abundant, an aliquot of a host culture available in our culture collection and parasites-free (Suppl. Table 2), was added to maximize the probability of an encounter between zoospores present in the sample and their hosts, and thus of new infections. When infected cells were observed under light microscopy, they were manually isolated and transferred to new wells to which dinoflagellate host culture was added, in order to establish a clonal parasitoid culture. The latter were further maintained by transferring a small volume to new wells and adding new healthy hosts twice a week.

### Single-cell PCRs from cultures

The cultured parasitoid strains were characterized at the molecular level by isolating individual mature sporangia using glass micropipettes. The sporangia were cleaned in several drops of autoclaved seawater, placed in 200-μL PCR tubes and stored at −80°C until processed. Those cells served as a template for single-cell PCRs in order to obtain 18S rDNA sequences, as described in (Reñé & Hoppenrath, 2019). Briefly, a PCR was conducted using primers EK82F – 28S-1611R (López-García et al., 2001; Moreira et al., 2007) to amplify the whole ribosomal operon. A subsequent PCR using the product of previous PCR as template and the primers EK82F – 1520R (López-García et al., 2001) was performed to amplify the 18S rDNA fragment. For cultures of the *Parvilucifera* genus, total genomic DNA was obtained from 10–15 mL of a culture containing both parasitoid and host. The culture was pelleted by centrifugation at 3,000 rpm for 15 min and the pellet then transferred to a 1.5-mL tube. After a second centrifugation, at 10,000 rpm for 5 min, the DNA was extracted using the DNeasy blood & tissue kit (Qiagen) following the manufacturer’s instructions. Based on a multiple sequence alignment of available Alveolata SSU rDNA sequences, ARB software (Ludwig et al., 2004) was used to design clade-specific Parv_F1 (5’- ATAGAGGCTTACCATGGC-3’) and Parv_R2 (5’-TCCACCTCTMACAAGCGA-3’) forward and reverse primers targeting the *Parvilucifera* clade, including *P. rostrata* (KF359483), *P. infectans* (KF359485, AF133909), *P. sinerae* (EU502912, KM878667) and *P. corolla* (KX519758, KX519761). The specificity of the PCR primers was checked in silico by submitting the sequences to the NCBI nr DNA (Primer-BLAST search) and SILVA (Testprimer search) databases (Pruesse et al., 2007; Ye et al., 2012). DNA amplification was performed by 25-μL reactions containing 1 μL of template DNA, 2.5 μL of 10× buffer, 1.5 mM MgCl_2_, 0.2 mM of each dNTP, 0.4 μM of forward and reverse primers, and 2 U of Taq DNA polymerase. The PCR program consisted of an initial denaturation of 94°C for 2 min, 35 cycles including a denaturation phase at 94°C for 15 sec, a polymerization phase at 55°C for 30 sec and an elongation phase at 72°C for 60 sec, followed by a final elongation phase at 72°C for 60 sec. Sanger sequencing was performed with both forward and reverse primers by an external service (Genoscreen, Lille, France).

### Preparation of mock communities and blank samples for metabarcoding

Mock microbial communities were processed using cultured strains available to serve as reference standards, thus ensuring correct amplification of the target organisms under the protocols used for metabarcoding. The cultures consisted of three dinoflagellate host strains belonging to *Alexandrium minutum*, *Ostreopsis* sp. and *Gymnodinium litoralis*, four Perkinsea strains belonging to *Parvilucifera sinerae*, *Tuberlatum* sp., *Dinovorax pyriformis* and Perkinsea sp. 1, and one strain belonging to the chytrid fungus *Dinomyces arenysensis*. All parasitoid strains were obtained during this study except the latter, which was previously available. Dinoflagellate strains were already available, as previously stated. Before the cultures were mixed, the cell abundances of the host and parasitoid strains were determined for further comparisons between the number of reads obtained from metabarcoding and the number of cells present in the samples. Thus, 1 mL of each strain was fixed in formaldehyde (1% final concentration) and the number of cells was counted using a Sedgewick Rafter chamber and Leica-Leitz DMII inverted microscope. For the parasitoid strains, both the infected cells (from early stages of infection to mature sporangium) and the healthy host cells present in the culture were counted. After the cell abundance of each culture was determined, 1 mL of each live strain was added to Falcon tubes in triplicate, resulting in 8 mL each. One replicate was serially filtered through polycarbonate filters as described above to serve as a positive control for procedures conducted with water samples; the other two replicates were gently mixed with 10 g of previously autoclaved sediment obtained from two sampling locations (Muga and Cubelles) and used as positive controls for the sediment samples. Blank samples (negative controls for the water samples) were prepared from 1 L of autoclaved MilliQ water serially filtered on 0.8- and 10-μm filters as previously described. All 0.8- and 10-μm filters and sediments containing the mock microbial communities and the blanks were stored at −80°C until further processing.

### DNA extraction and metabarcoding sequencing

Total genomic DNA was extracted from all filters, including all environmental water samples, mock communities and blank samples, using the DNeasy PowerSoil kit (Qiagen) as follows: Filters cut in small pieces using sterilized razor blades were placed in a PowerBead to which 60 μL of solution C1 was added. The tubes were vortexed at maximum speed for 10 min and then centrifuged at 10,000 x*g* for 30 s. The supernatant was transferred to a clean collection tube, removing all filter pieces. Successive steps were conducted according to the manufacturer’s instructions. For sediment samples, including environmental and mock communities, total genomic DNA was extracted using the DNeasy PowerMax soil kit (Qiagen) and 10 g of sediment sample, following the manufacturer’s instructions. DNA was eluted in 5 mL and concentrated to a final volume of 200 μL using Amicon Ultra centrifugal filters. A first PCR was conducted using universal eukaryotic primers designed against metazoan EK-565F (5’-GCAGTTAAAAAGCTCGTAGT-3’) (Simon et al., 2015) and 18S-EUK-1134-UnonMet (5’-TTTAAGTTTCAGCCTTGCG) (Bower et al., 2004), resulting in an amplicon length of around 560 base pairs. DNA amplification was performed by 25-μL PCR reactions containing 2 μL of template DNA, 2.5 μL of 10× buffer, 1.5 mM MgCl2, 0.2 mM of each dNTP, 0.4 μM of forward and reverse primers, and 2 U of Taq DNA polymerase. The PCR program consisted of an initial denaturation at 94°C for 2 min, 35 cycles including a denaturation phase at 94°C for 15 sec, a polymerization phase at 55°C for 30 sec and an elongation phase at 72°C for 60 sec, followed by a final elongation phase at 72°C for 60 sec. Four μL of each product was electrophoresed in a 1.2% agarose gel stained with SYBR Safe (Invitrogen) for visualization of the bands and confirmation of the correct amplicon length. The remaining PCR product was purified using the QIAquick PCR purification kit (Qiagen) and sent to an external service (AllGenetics, A Coruña, Spain), where a semi-nested PCR using primers with the adaptors EK-565F and E1009R (5’- AYGGTATCTRATCRTCTTYG-3’) (Comeau et al. 2011) was performed, followed by MID-indexing of the products, their pooling in equimolar amounts and sequencing in a fraction (1/2) of a MiSeq PE300 Illumina run.

### Molecular data processing

The raw reads obtained from previous samples were trimmed from the adaptors using cutadapt v.1.16 (Martin, 2011). The DADA2 pipeline (Callahan et al., 2016) was used to filter, dereplicate and trim low-quality sequences (maxEE = 4, 6; truncLen= 270, 250), merge the paired ends (minOverlap = 10) and remove chimeras. The resulting ASVs (amplicon sequence variants) were classified using a simple non-Bayesian taxonomy classifier, vsearch v2.8.1 (Rognes et al., 2016), and the PR2 database (Guillou et al., 2013) customized with new in-house Perkinsea sequences; the bootstrap cut-off was 0.6. Raw metagenomic reads were submitted to the NCBI Sequence Read Archive under the bioproject number PRJNA630546.

### Phylogenetic analysis

Partial 18S rDNA sequences obtained from cultured strains and all ASVs belonging to Perkinsea obtained from the environmental samples were aligned with a selection of sequences obtained from GenBank representing the phylogenetic diversity of Perkinsea. All reference sequences were aligned using the auto option of MAFFT online v.7. The ASVs were then included in the backbone alignment using the add option of MAFFT online v.7. The resulting alignment was trimmed using TrimaL, resulting in a final alignment of 1840 positions. Phylogenetic relationships were determined using maximum-likelihood (ML) and Bayesian inference methods. For the former, the GTRGAMMA evolution model was used on RAxML v. 8.2. All model parameters were estimated using RAxML. Repeated runs on distinct starting trees were carried out to select the tree with the best topology (the one with the greatest likelihood of 1000 alternative trees). The bootstrap ML analysis was done with 1000 pseudo-replicates and the consensus tree was computed with the RAxML software. Bayesian inference was performed using MrBayes v.3.2, run with a GTR model in which the rates were set to gamma. Each analysis was performed using four Markov chains, with one million cycles for each chain. The consensus tree was created from post-burn-in trees and the Bayesian posterior probabilities of each clade were examined.

### Statistical analysis

The metabarcoding data were analysed using the package phyloseq (McMurdie & Holmes, 2013), from Rcran, and all graphs were generated using the package ggplot2 (Wickham, 2016). The saturation of the samples was evaluated in rarefaction curve plots using the package vegan (Oksanen et al. 2020), to determine whether the sequencing effort was sufficient to capture most of the diversity. Nonmetric multidimensional scaling (NMDS) using Bray-Curtis distances on Hellinger-transformed counts for all ASVs was then performed to analyse the dissimilarity between samples and replicates, including the blanks and mock communities that served as negative and positive controls, respectively. The results are shown in Suppl. File, Suppl. Figs. 2–4. The ASVs belonging to Dinophyceae were then subsampled and used to evaluate both the dinoflagellate community and the dissimilarity between samples based on the same parameters mentioned above, to visualize the previously determined groups of blooming host species. The significance of the observed groups was assessed in an analysis of similarities (ANOSIM) (Clarke et al. 1993) from the package vegan. Both a global test and pairwise tests by bloom group were performed, applying Bray-Curtis distances with 999 permutations and a significance cut-off for the p-value of 0.05. Heatmap visualization and clustering were done using the Hellinger-transformed counts of the ASVs belonging to Perkinsea. The dendrogram was generated using average linkage clustering with Euclidean distances; the plot was created using the heatmap.3 package (Griffith 2018). The significance of the different groups obtained from the Perkinsea community was also statistically tested in an ANOSIM as previously described. To further identify taxa indicative of a particular blooming host group, an indicator species analysis, which evaluates the specificity and fidelity of each species for the different groups, was performed employing the IndVal function in the labdsv package (Roberts 2019). Only those ASVs with IndVal values ≥ 0.5 and p-values < 0.05 were retained.

## RESULTS

### Diversity and abundance of the dinoflagellate community

The number of eukaryotic reads obtained from the 101 samples ranged from 6,158 to 55,217 (Suppl. Table 1), with each sample showing richness saturation (Suppl. Fig. 1). The number of ASVs ranged from 17 to 399. Reads belonging to dinoflagellates (class Dinophyceae) were detected in all samples, in proportions of 0.6–99.8%. Dinoflagellates were the most highly represented (mean 74.9% ± 24.2) in the water column samples from the large size fraction (10–200 μm), while lower relative abundances were retrieved from the sediment samples (mean 24.1% ± 19.7). The number of ASVs belonging to dinoflagellates ranged from 3 to 64. As previously described, samplings were performed when target species abundances >10^4^ cells L^−1^ were detected by microscopy analysis. In most cases, those species were predominant in the community, representing 60-90% of dinoflagellate individuals. However, this was not always the case. In L’Estartit, the dominant species was *Prorocentrum triestinum* (Fig. 2); in Fra Ramon, *K. foliaceum* represented approx. 41% of the total dinoflagellate community and *Ansanella* sp. 51%. The correspondence between microscopy counts and metabarcoding sequencing was also explored (Fig. 2), with the latter based on the metabarcoding results from the large-fraction samples (>10 μm), as they better represented the dinoflagellate community. For samples obtained during *A. minutum* blooms, there was good agreement except in the case of L’Estartit, in which the representation of *A. minutum* was higher by metabarcoding than by microscopy. For samples corresponding to *A. taylorii* and *G. litoralis* blooms, a high level of agreement between the two techniques was determined except at Aiguafreda. Samples corresponding to *Ostreopsis* sp. blooms were dominated by this species based on the microscopy results and those of metabarcoding, but with a smaller contribution in the latter. Finally, at Fra Ramon, *K. foliaceum* was not the dominant species in terms of cell abundance, and its proportion was smaller in metabarcoding results than in microscopy counts. The samples obtained during no-bloom conditions of the target species showed a heterogeneous species composition. The differences between the two methodologies were mostly related to the presence in the metabarcoding analyses of the small dinoflagellate *Ansanella* sp. which could not be identified by microscopy.

**Figure 2:**
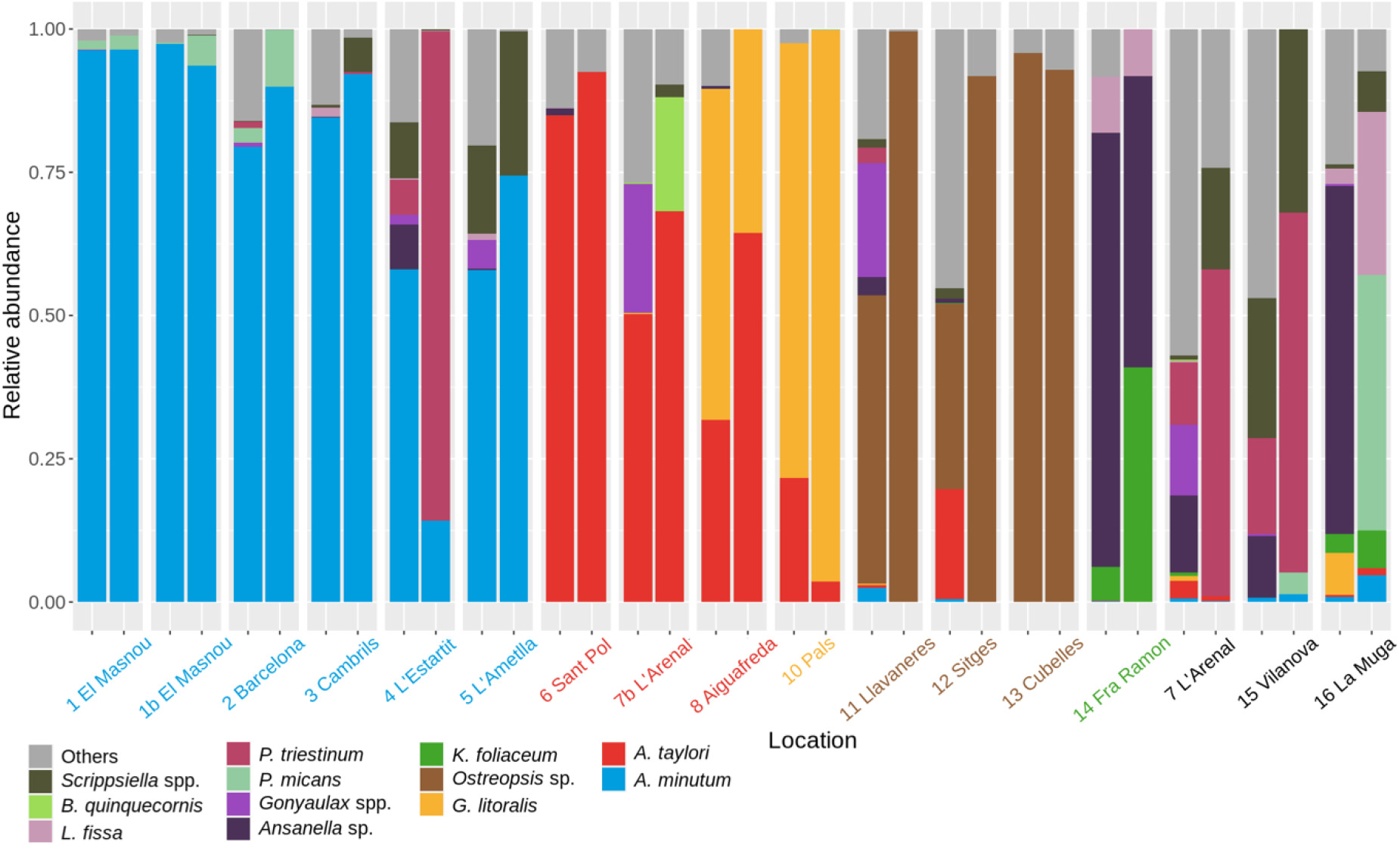
Barplots showing the relative abundance of each dinoflagellate species detected in the studied samples. For each location, two bars are shown. The first one represents the metabarcoding results and the second one the microscopy results. Samples are grouped based on the target dinoflagellate species (colors from X axis). Black ones correspond to those sampled during no-bloom conditions of previous target species.

### Diversity of Perkinsea parasitoids along the Catalan coast

Reads belonging to the class Perkinsea were detected in 15 of the 19 sampled locations, corresponding to 40 of the 101 samples, nine of them belonging to samples from the small size fraction, 12 to the large fraction and 19 to the sediments. However, in nine additional samples belonging to the large water-column fraction (sample sites of El Masnou, Aiguafreda, L’Ametlla de Mar and Pals) the relative abundance of dinoflagellate reads was >94%, such that other community components were scarcely represented and Perkinsea reads were not detectable (Suppl. Table 1). The relative abundance of Perkinsea was always <1%, except in three sediment samples from Cubelles, in which Perkinsea reads represented ~30% of the relative abundance of the eukaryotic community. No more than five Perkinsea ASVs were detected in the same sample.

The combination of cultivation and metabarcoding allowed a detailed identification of the Perkinsea species infecting the dinoflagellates present in the sampling locations and a determination of the host infected by those species in nature. Infected host cells were observed in incubations performed with field samples, which enabled the establishment of eight culture strains belonging to four different Perkinsea species and their subsequent identification and molecular characterization (Fig. 3, Suppl. Table 2). Additionally, metabarcoding results allowed the recovery of 32 ASVs attributed to the class Perkinsea (Fig. 4). Those reads were detected in all sampling locations except El Masnou Harbour, Vilanova i la Geltrú Harbour and the beach at Aiguafreda. Seventeen ASVs, some representing different haplotypes for the same species, could be directly related to nine different Perkinsea infecting dinoflagellates, including seven that corresponded to previously known species and two new species yet to be described. Eight additional ASVs could not be assigned to a given species, but they clustered close to Parviluciferaceae and likely belonged to as-yet-undescribed species infecting dinoflagellates. The remaining seven ASVs did not seem to be closely related to previous parasitoid species and cannot be assumed to correspond to dinoflagellate parasitoids. More than one Perkinsea ASV co-occurred in 19 of 40 samples, demonstrating the co-existence of different parasitoids of dinoflagellates at the same sampling location (Suppl. Table 1). All eight cultivation-based detections were also detected by metabarcoding (Table 2), with the exception of *Parvilucifera sinerae* from El Masnou Harbour. In the case of Perkinsea sp. 1, representing a species not yet described, it could be identified in the metabarcoding dataset after its 18S rDNA sequence was obtained from the culture. The remaining 18 detections were limited to the metabarcoding approach and for *Parvilucifera rostrata, P. catillosa* or *Tuberlatum coatsi* constituted the first reported detections in the studied area.

**Table 2:**
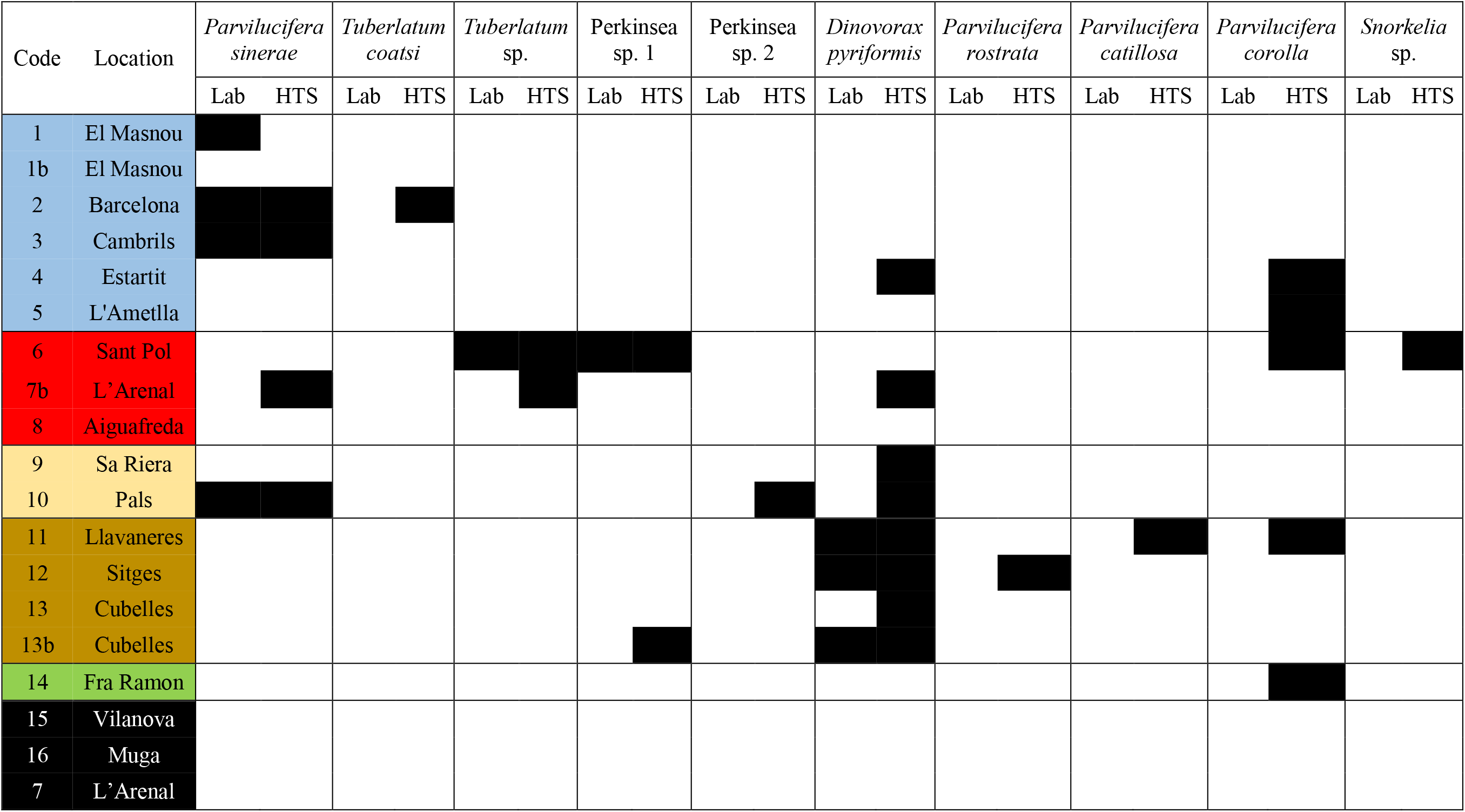
Comparison of dinoflagellate parasitoids detected during this study in the laboratory by cultivation (Lab) and by metabarcoding (HTS) for each sampling. The colours indicate the bloom-forming dinoflagellate species present in the sample: *Alexandrium minutum* (blue), *A. taylorii* (red), *Ostreopsis* sp. (brown), *Gymnodinium litoralis* (yellow), *Kryptoperidinium foliaceum* (green), and no-bloom of target host (black).

**Figure 3.**
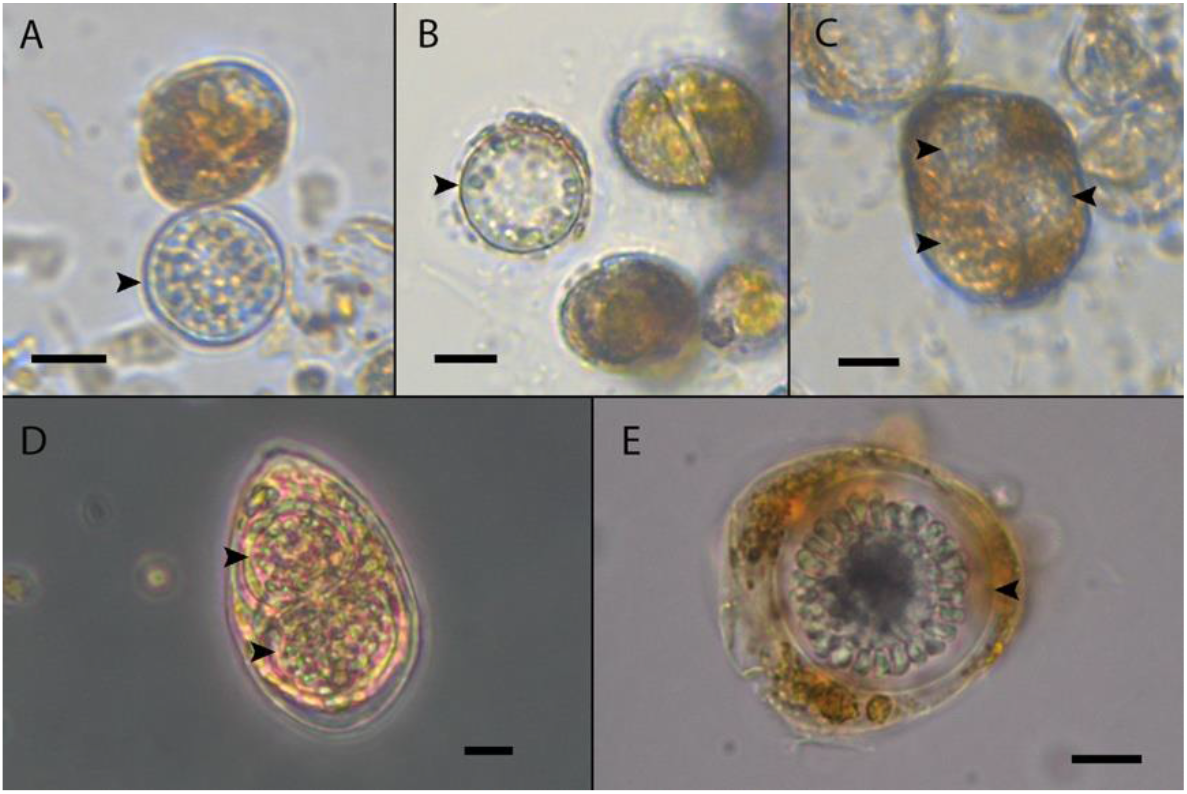
Light microscopy images of Perkinsea infections (arrowheads) on dinoflagellates. A) Early sporangium of *Parvilucifera sinerae* infecting *Alexandrium minutum*. B) Late trophont of *P. sinerae* infecting *Gymnodinium litoralis*. C) Triple infection by *Tuberlatum* sp. of *Alexandrium taylorii*. D) Double infection by *Dinovorax pyriformis* of *Ostreopsis* sp. E) Late sporangium of Perkinsea sp. 1 infecting *A. taylorii*. Scale bars = 10 μm.

**Figure 4.**
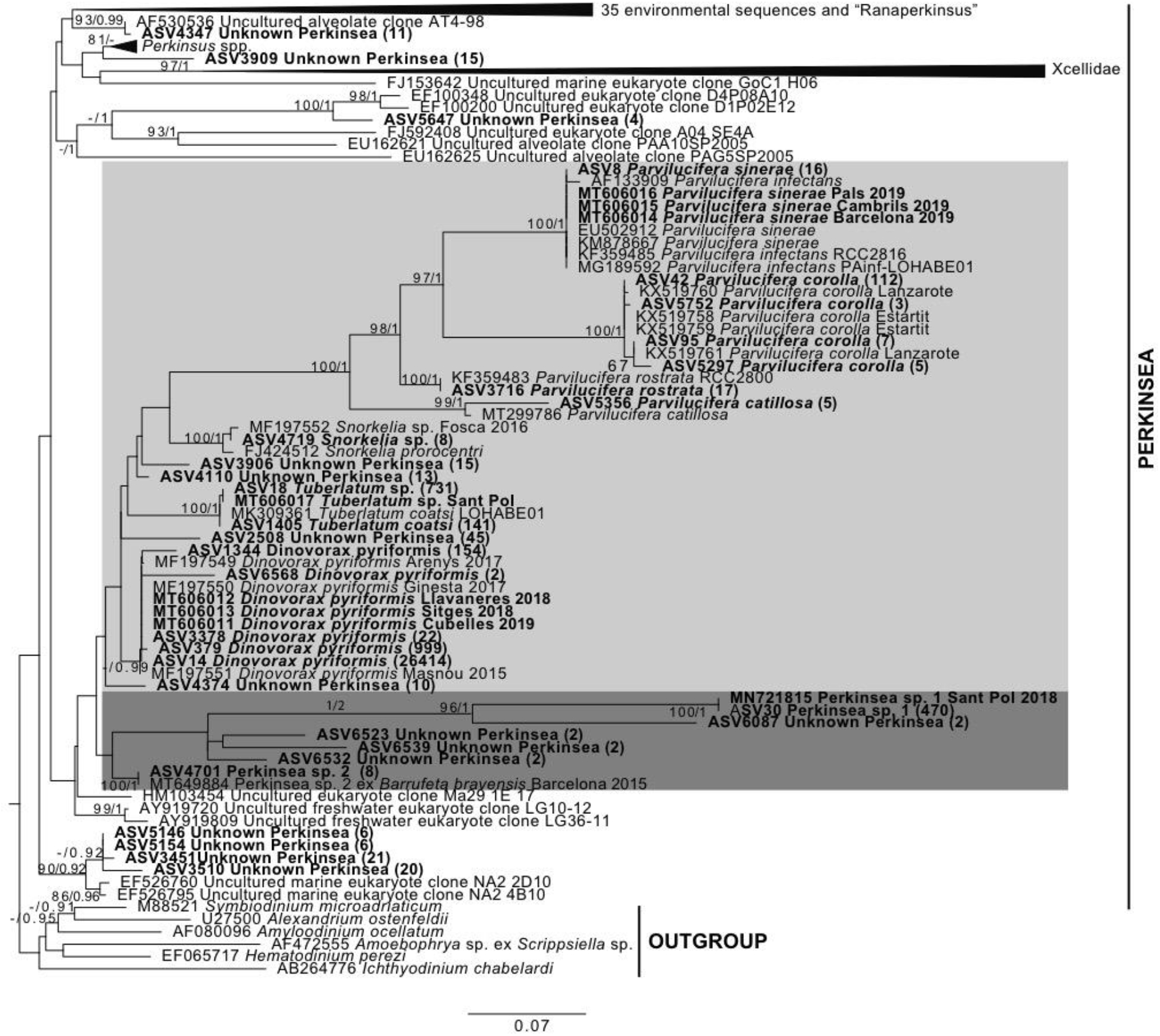
Maximum-likelihood phylogenetic tree of 18S rDNA. Reference sequences of Perkinsea representatives and ASV sequences of V4 18S rDNA from this study are shown in bold. Dinoflagellate and MALV representatives served as outgroups. The numbers in parentheses after the ASV label indicate the total number of reads, and the numbers in nodes the bootstrap values (%) and Bayesian posterior probability. The area in light grey includes Parviluciferaceae members, and that in dark grey other parasitoids of dinoflagellates.

### Determination of parasite-host interactions

The NMDS analysis using the metabarcoding data of the dinoflagellate community resulted in a robust categorization of the samples according to their blooming host species (Fig. 5). Thus, samples corresponding to five categories (*A. minutum*, *A. taylorii*, *G. litoralis*, *K. foliaceum* and *Ostreopsis* sp.) contained distinctive dinoflagellate communities. Some additional sampling locations were sampled during no-bloom conditions. Blooms of *A. minutum* are recurrent at Vilanova i la Geltrú Harbour, and despite the low abundance of this species during sampling conducted, the samples clustered within the *A. minutum* category. La Muga and L’Arenal clustered together but separately from the other four categories, while their sediment samples clustered within the category *Ostreopsis* sp. The ANOSIM analysis based on the Bray-Curtis distance matrix revealed significant differences in the dinoflagellate community between the different groups (Table 3), allowing further exploration of the differences in their parasitoid community. Pairwise tests showed statistically significant differences between specific groups. However, the R values were generally in the intermediate range (0.25-0.5), suggesting some degree of overlap between them, as shown in Fig. 5.

**Table 3:** Results of the global and pairwise tests obtained from the analysis of similarity (ANOSIM) conducted between the different host groups established, based on the dinoflagellate community. Tests showing statistical significance (p<0.005) are in bold.

|  |  |  |  |  |  |  |
| --- | --- | --- | --- | --- | --- | --- |
| Global test<br>**R=0.392;**<br>**p=0.001** | *A. minutum* | *A. taylori* | *G. litoralis* | *K. foliaceum* | *Ostreopsis* sp. | No-bloom |
| *A. minutum* |  |  |  |  |  |  |
| *A. taylori* | **R=0.284;**<br>**p=0.001** |  |  |  |  |  |
| *G. litoralis* | **R=0.25;**<br>**p=0.001** | **R=0.186;**<br>**p=0.001** |  |  |  |  |
| *K. foliaceum* | **R=0.201;**<br>**p=0.001** | **R=0.264;**<br>**p=0.001** | **R=0.551;**<br>**p=0.002** |  |  |  |
| *Ostreopsis* sp. | **R=0.301;**<br>**p=0.001** | **R=0.316;**<br>**p=0.001** | **R=0.402**<br>**p=0.001** | **R=0.334;**<br>**p=0.001** |  |  |
| No-bloom | **R=0.155;**<br>**p=0.001** | **R=0.101;**<br>**p=0.001** | **R=0.254;**<br>**p=0.001** | **R=0.147;**<br>**p=0.001** | **R=0.249;**<br>**p=0.001** |  |

**Figure 5.**
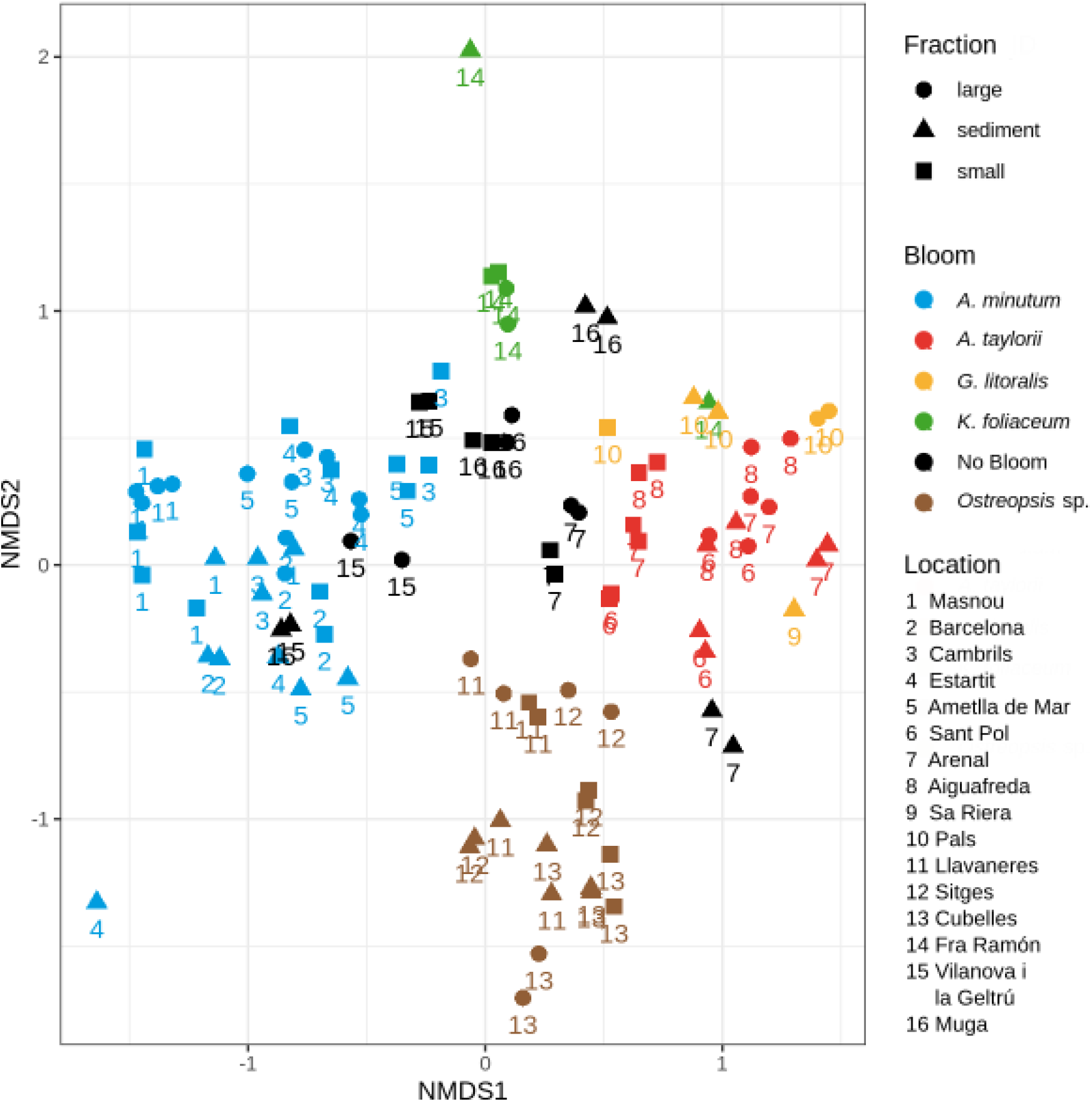
Non-metric multidimensional scaling (NMDS) ordination plot, based on Bray-Curtis distances for the dinoflagellate community. The different colours indicate the different blooming species, and the different shapes the sample fraction (small, large or sediment). Labels correspond to the sampling locations and are explained in Table 1 and Figure 1.

The Perkinsea community was evaluated based on the samples with positive detections, i.e. 40 of the 101 samples obtained. The different host categories were dominated by different parasitoid species. Some species, such as *Tuberlatum coatsi*, *Tuberlatum* sp., and Perkinsea sp. 1, exclusively appeared in a specific dinoflagellate category (Fig. 6), and others, such as *Dinovorax pyriformis*, *Parvilucifera corolla*, and *P. sinerae*, appeared in most categories. Moreover, their relative abundances varied depending on the causative blooming species. The samples were grouped based on the similarity of the Perkinsea community (Fig. 7), yielding clearly defined clusters. The first cluster included samples related to the *Dinovorax pyriformis* detections, mainly corresponding to samples representing *Ostreopsis* sp. and *G. litoralis* blooms. The second cluster contained samples with *P. corolla* detections, corresponding to those from *K. foliaceum* blooms, but also blooms of *A. minutum* and *Ostreopsis* sp. The third cluster comprised detections of *Tuberlatum* sp. and Perkinsea sp.1, corresponding to samples from sites of *A. taylorii* blooms. A forth cluster was characterized by the presence of *T. coatsi* and *P. sinerae*, comprising samples affected by *A. minutum* blooms. Samples obtained from two locations during no-bloom conditions (Muga and L’Arenal) clustered independently, but both locations were characterized by the presence of ‘unknown Perkinsea’. Finally, there were also samples that were not assigned to any of these clusters. For example, only one Perkinsea species (Perkinsea sp. 2, *Parvilucifera catillosa*, and *D. pyriformis*) was detected in each of the samples from three different categories (*G. litoralis*, *Ostreopsis* sp., and *A. minutum* respectively), and they lacked similarity with the rest of the samples from their corresponding category. An ANOSIM test was conducted to evaluate the statistical significance of the Perkinsea community determined for each host group (Table 4). Statistically significant differences between all groups were determined in the global test and in most pairwise tests conducted between all specific groups, especially the communities found during *A. taylorii* and *Ostreopsis* sp. blooms. However, parasitoid communities from the *G. litoralis* blooms significantly (p<0.005) differed only when compared to those from *A. taylorii*, and the parasitoid communities from *K. foliaceum* blooms significantly differed only when compared to those from *A. taylorii* and *Ostreopsis* sp. blooms. An analysis of indicator ASVs was performed to test the statistical significance of those observations by delineating high fidelity differential abundance patterns (IndVal values ≥ 0.5, p < 0.01). *Parvilucifera corolla* was the top scoring taxon for *K. foliaceum* blooms (IndVal=0.79, p=0.002); *Tuberlatum* sp. and Perkinsea sp.1 for *A. taylorii* blooms (IndVal=0.89, p=0.001; IndVal=0.67, p=0.016 respectively), and *T. coatsi* for *A. minutum* blooms (IndVal=0.55, p=0.03). Samples obtained during no-bloom conditions were related to the presence of different unknown Perkinsea species (IndVal>0.50, p<0.05). *Dinovorax pyriformis* was associated with *Ostreopsis* sp. blooms but although the association was statistically significant, the IndVal value did not reach the threshold (IndVal=0.48, p=0.012).

**Table 4:** Results of the global and pairwise tests obtained from the analysis of similarity (ANOSIM) conducted between the different host groups established, based on the Perkinsea community. Tests showing statistical significance (p<0.005) are in bold.

|  |  |  |  |  |  |  |
| --- | --- | --- | --- | --- | --- | --- |
| Global test<br>**R=0.567;**<br>**p=0.001** | *A. minutum* | *A. taylorii* | *G. litoralis* | *K. foliaceum* | *Ostreopsis* sp. | No-bloom |
| *A. minutum* |  |  |  |  |  |  |
| *A. taylorii* | **R=0.542;**<br>**p=0.001** |  |  |  |  |  |
| *G. litoralis* | R=0.266;<br>p=0.077 | **R=0.779;**<br>**p=0.005** |  |  |  |  |
| *K. foliaceum* | R=0.121;<br>p=0.226 | **R=0.918;**<br>**p=0.022** | R=0.5;<br>p=0.2 |  |  |  |
| *Ostreopsis* sp. | **R=0.558;**<br>**p=0.001** | **R=0.774;**<br>**p=0.001** | R=0.151;<br>p=0.261 | **R=0.75;**<br>**p=0.015** |  |  |
| No-bloom | **R=0.327;**<br>**p=0.004** | **R=0.833;**<br>**p=0.002** | R=0.333;<br>p=0.105 | R=0.428;<br>p=0.2 | **R=0.77;**<br>**p=0.002** |  |

**Figure 6.**
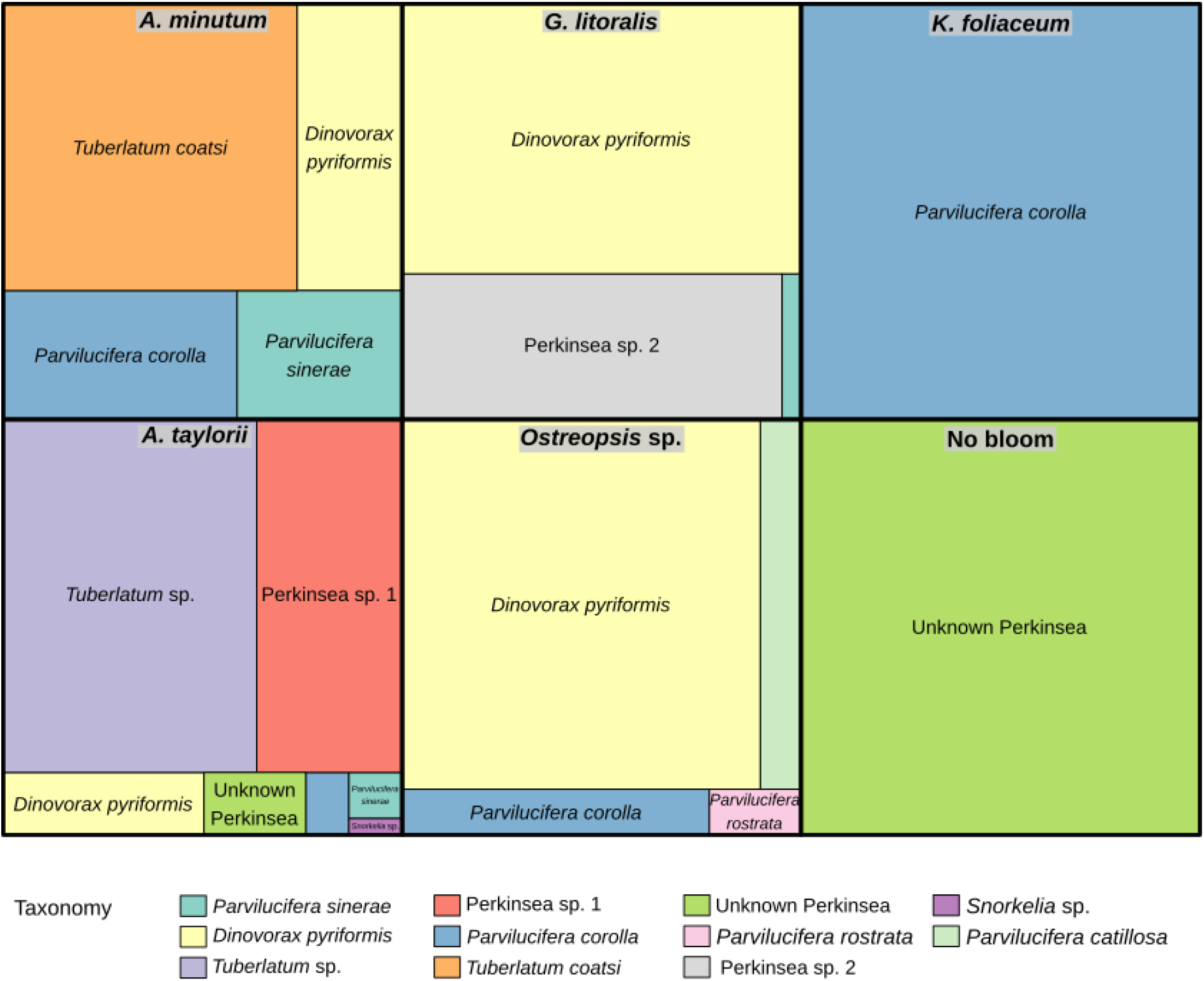
Relative abundance of ASVs belonging to Perkinsea obtained in all samples from each studied dinoflagellate bloom. All ASVs belonging to the same species were agglomerated. In the case of ‘Unknown Perkinsea’, the boxes correspond to five different ASVs/species in *A. taylorii* and seven in the no-bloom category. Only species with a relative abundance > 0.5% are shown.

**Figure 7.**
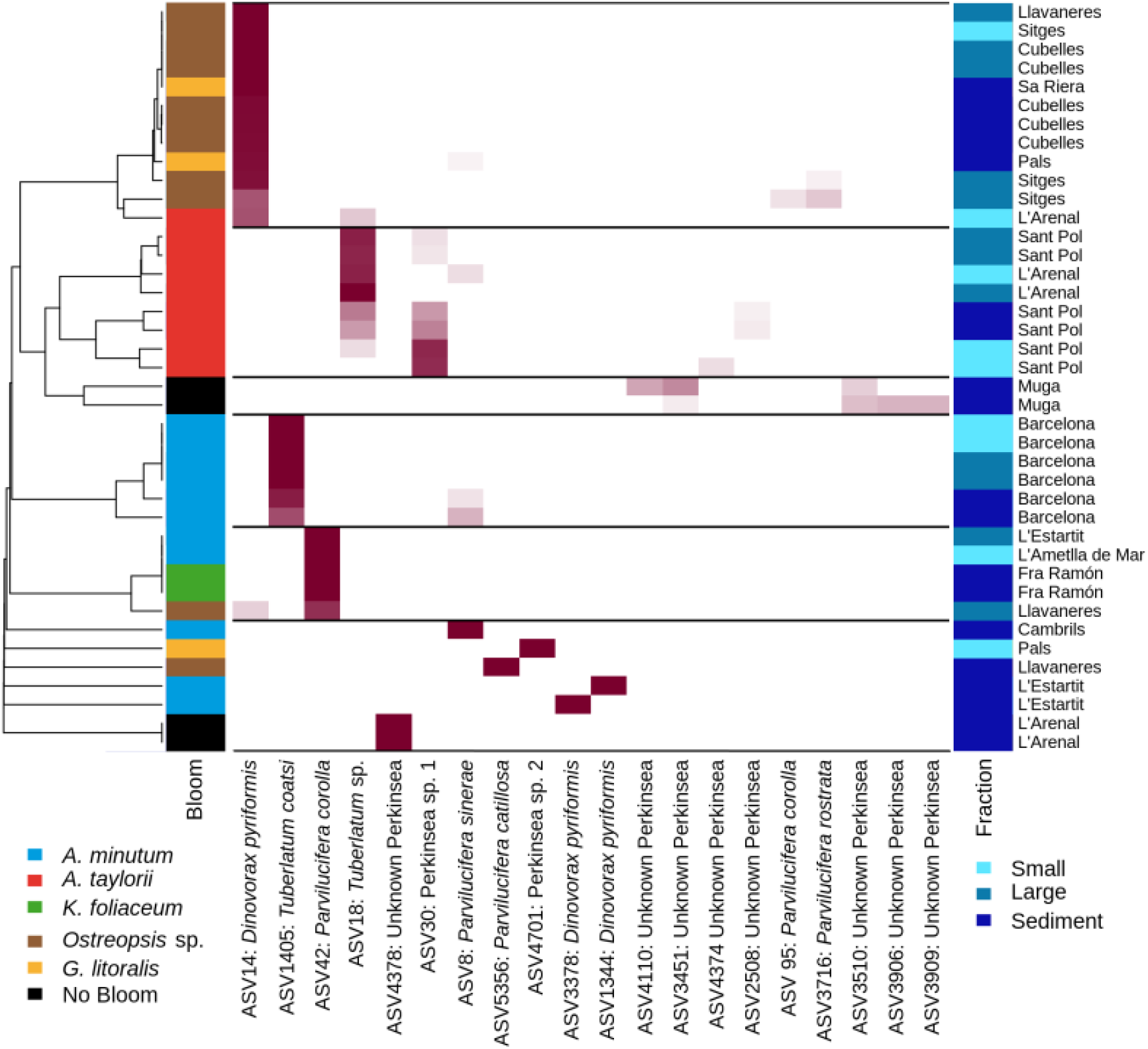
Heatmap of the 20 most abundant Perkinsea ASVs (columns) identified in all samples (rows). Annotation bars beside the heatmap report the fraction (right) and bloom group (left). The dendrogram was generated using average linkage clustering with Euclidean distances based on the relative abundances of the taxa in each sample.

## DISCUSSION

### Diversity and distribution

Parasites of protists have been studied for decades by direct observations, which have provided knowledge on chytrids (basal fungi) (Canter, 1984) and stramenopiles (Drebes, 1966; Kühn et al., 1996). Following the application of genetic tools, the molecular diversity of parasitic groups has been examined in depth by environmental sequencing, which allows determinations of parasite distributions based on molecular information obtained from discrete observations (Alves-de-Souza et al., 2019; Garvetto et al., 2019; Garvetto et al., 2018) as well as the development of specific molecular probes to investigate parasitic interactions between protists (Chambouvet et al., 2019). However, few studies have combined a molecular approach with a direct characterization from cultures and field samples to determine the diversity and ecology of a parasitic group. Among the studies of Perkinsea species belonging to Parviluciferaceae (parasitoids of dinoflagellates), many consist of discrete observations made at specific locations. Thus, while the presence of *Parvilucifera infectans* / *P. sinerae* has been widely reported (Jeon et al., 2018), other species such as *P. rostrata* are known only from the French Atlantic coast (Blanquart et al., 2016; Lepelletier et al., 2014), *P. catillosa* has thus far been described only from the Baltic Sea (Alacid et al. 2020), and the genus *Tuberlatum* and *P. multicavata* only from Korean waters (Jeon & Park, 2019; Jeon & Park, 2020). In this study, a comparison of the different tools used was evaluated to check the reliability of the knowledge obtained. Metabarcoding results showing the dominance of bloom-forming dinoflagellate species were consistent with microscopy observations. Although discrepancies between the two methods in the relative abundances of the target species were also detected, they did not alter our definition of a bloom as a cellular abundance >10^4^ cells L^−1^ and blooms of the different dinoflagellate communities allowed a determination of the parasitoid species that predominated under the different respective conditions. The overall diversity and distribution of the Perkinsea community at the study area determined by combining morphological and molecular characterizations led to the detection not only of many known species, but also of new species, as yet undescribed. The specific host-parasitoid interactions could be determined *in situ* based on direct observation of field samples and the establishment of cultures. However, because not all parasitoid species can be grown in the laboratory, metabarcoding was performed as well, and almost all established host-parasitoid co-cultures were also identified by metabarcoding.

Most known Parviluciferaceae species were detected in our survey of the Catalan coast, such that our results can be considered as representative of Perkinsea diversity in the study area. Additionally, many of the obtained ASVs clustered close to other known species and likely corresponded to unknown dinoflagellate parasitoids, thus highlighting the need for further investigations of parasitoid diversity and ecology. By contrast, none of the ASVs of unknown affiliation detected under no-bloom conditions could be assigned to dinoflagellate parasitoids with the exception of ASV4110 detected in sediments from La Muga. This result demonstrates that dinoflagellate blooms offer optimal conditions for the growth and propagation of parasitoids, as in their absence, parasitoids are not detected. Two hypotheses can be proposed to explain parasitoid survival and maintenance in the absence of a bloom: (1) Parasitoids could infect non-preferred hosts at very low-prevalence levels, which would ensure their survival until preferred host abundances are high enough to support propagation. (2) Like their hosts, the parasitoids may enter a latent phase in the sediments until favourable conditions return, as demonstrated for *Amoebophrya* parasitoids (Chambouvet et al., 2011). The detection in our study area of many parasitoid species known only from distant locations is indicative of the widespread distribution of this group, consistent with the global distribution of their hosts. Finally, the distribution of many parasitoid species at the Catalan coast was not restricted to a specific location, but was in agreement with that of their preferred hosts.

### Host preferences in parasite-host interactions

Previous laboratory experiments examining the host range of Perkinsea revealed the ability of the parasitoids to infect a large number of dinoflagellate hosts, suggesting that they were generalists. However, differences in host susceptibility to infection, even at the intraspecific level, have been described (Garcés et al., 2013; Lepelletier et al., 2014; Rodríguez & Figueroa, 2020). Additional experiments demonstrated that despite their potential to infect multiple species, the parasitoids exhibited clear host preferences, selecting one host over another, equally available one (Alacid et al., 2016). Field studies have explored parasitoid-host interactions and dynamics. In two estuaries in France, the strong correlation between the presence of *A. minutum* and the parasitoids *P. infectans* and *P. rostrata* suggested that parasitoids are dependent on the presence of their preferred host at high abundances (Blanquart et al. 2016). In another study, conducted at a harbour in the Mediterranean Sea, the dynamics of *P. sinerae* were directly linked to *A. minutum* bloom development (Alacid et al. 2017). Although our study did not evaluate the succession of hosts and parasites at each location to determine the changes occurring over time, we did explore the parasitoid-host interactions occurring along a defined geographical area characterized by recurrent blooms of several dinoflagellate species, of which five were included in this study (*A. minutum*, *A. taylorii, K. foliaceum, G. litoralis,* and *Ostreopsis* sp.), and by no-bloom periods. Our results demonstrated both the dependency and preferences of parasitoid communities on a blooming host. Thus, while a parasitoid is able to infect a wide range of dinoflagellate species, our investigation suggests that a bloom of the preferred host provides optimal conditions for propagation. This conclusion is supported by our additional finding of the absence of the parasitoids in locations where the abundance of its host during sampling was low. Further investigation of parasitoid preferences revealed different parasitoid species at the same location, as shown in Fig. 6 and Fig. 7. Different parasitoids co-occurred in the same location and infected the same host, e.g. *Tuberlatum* sp. and Perkinsea sp. 1, implying some degree of competition or cooperation between them, an observation that merits further study. In other cases, different species were detected but only one of them was dominant. Such observations are consistent with previous knowledge of parasitoid host-ranges and host preferences. For instance, blooms of *A. minutum* are recurrent at Arenys de Mar harbour and *P. sinerae* is often found in association with them (Figueroa et al., 2008; Alacid et al. 2017). Following the decay of these blooms, the dinoflagellate *Prorocentrum micans* appears in high abundances and serves as a host for infection by *D. pyriformis* (Reñé et al. 2017b).

Dinoflagellate community succession implies a change in the dominant species at a given location, as the niche vacated by one species becomes occupied by another. In turn, this allows the proliferation of the parasitoid species with a preference for that host (Fig. 8). Additionally, host preferences are not restricted to a single host species, and infections of diverse hosts in nature by a single parasitoid species are known from the literature. For instance, the above-mentioned *D. pyriformis* was the dominant species in samples with high abundances of *Ostreopsis* sp. and in the locations where *Prorocentrum* and *Dinophysis* species were present, a coupling described in a previous report (Reñé et al., 2017b). Similarly, *P. sinerae* is commonly detected during blooms of *A. minutum* in the Catalan coast, predominantly in harbours (Turon et al., 2015), but in this study it was also observed infecting *G. litoralis*, which proliferates in beaches, and its preference for *Scrippsiella trochoidea* was previously demonstrated (Alacid et al. 2016). Thus, consistent with laboratory demonstrations of their wide host range, Perkinsea parasitoids are able to infect different host species in nature, regardless of the geographical areas where proliferate. In summary, while parasitoids have a direct impact on the growth and abundance of dinoflagellates, our results suggest that dinoflagellates shape the parasitoid community, by representing their ecological niche and influencing parasitoid abundance and distribution.

**Figure 8.**
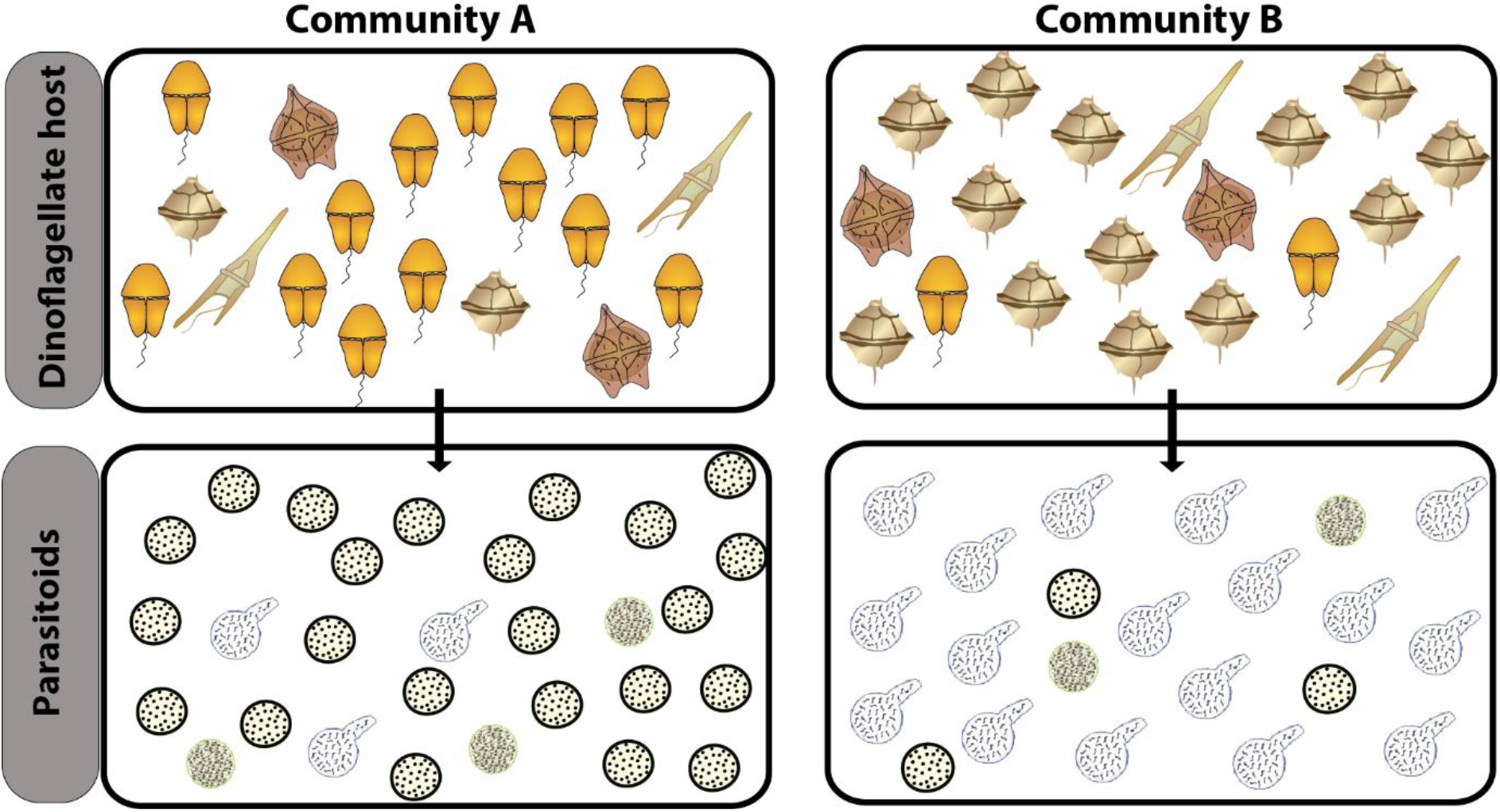
Schematic drawing showing two dinoflagellate blooms situations. For community A, the abundances of the parasitoid A increase when its preferred dinoflagellate host A is present at high abundances. For community B, the parasitoid B increases its abundances when its preferred dinoflagellate host B is predominant.

## CONCLUSIONS

1. Dinoflagellate blooms provide optimal conditions for the propagation of parasitoid infections.
2. Almost all known species of Perkinsea infecting dinoflagellates are present at the Catalan coast, suggesting a wide geographical distribution of these parasitoids. Some species detected have yet to be described.
3. The widespread distribution of Perkinsea species follows that of their respective host(s), which may result in the coexistence of different parasitoid species in the same geographic location.
4. The specific parasite-host interactions determined for each of the studied blooms demonstrated the host preferences exhibited by parasitoids in nature. The dominance of a species within the parasitoid community is driven by the presence and abundances of its preferred host(s).

## ACKNOWLEDGEMENTS

This study was funded by the MINECO Project COPAS ‘Understanding top-down control in coastal bloom-forming protists’ (CTM2017-86121-R) and the institutional support of the ‘Severo Ochoa Centre of Excellence’ accreditation (CEX2019-000928-S). We thank A. Chambouvet (CNRS, France) for designing the *Parvilucifera* specific primers. E. Alacid is funded by the Royal Society through a Newton International Fellowship. A.D. Fernández-Valero is funded by the MICIU grant “Ayudas para contratos predoctorales para la formación de doctores 2018 (PRE2018-084893)”

## DATA ACCESSIBILITY

DNA sequences from cultures are available in GenBank, accessions MT606011 – MT606017, MN721815; metabarcoding raw sequences and all samples metadata are available in NCBI SRA PRJNA630546.

## AUTHOR CONTRIBUTIONS

AR and EG designed the study; all authors performed samplings and/or laboratory procedures; AR, NT, NS, JG and EG analysed data; AR wrote the manuscript with the contribution of all coauthors.

## Notes

### Competing Interest Statement

The authors have declared no competing interest.

