## Supplementary Figure 1, Figure 2, Figure 3, Figure 4, Table 1 and Table 2 for "Host preferences of coexisting Perkinsea parasitoids during coastal dinoflagellate blooms"

### Evaluation of blank and mock samples

Negative (blank) and positive (mock) controls were processed during metabarcoding sequencing as reference standards. One blank sample containing autoclaved MilliQ water and one mock sample containing dinoflagellate and parasitoid cultured cells at known abundances were processed by serial filtration, yielding one small (0.8–10 µm) and one large (10–200 µm) fraction for each sample. Two samples of autoclaved sediments obtained from different locations containing the mock community were also processed.

All samples analyzed showed richness saturation (Suppl. Fig. 1), indicating that the sequencing effort was sufficient to obtain a robust characterization of the microeukaryotic community. An MDS analysis was then performed with the community information to evaluate the similarity between locations, fractions (small, large and sediment) and sample replicates (Suppl. Fig. 2). Blank and mock samples were also included in the analysis to use them as negative and positive controls, respectively. All samples corresponding to the same location generally clustered together, even though the similarity between small and large seawater fractions was higher than for sediment ones. All sample replicates were in agreement, showing high similarity.

The results from the two fractions of blank sample differed (Suppl. Fig. 2). The small fraction contained only a few ASVs with a low number of reads, and its community composition was not related to that of the study samples. By contrast, the community composition of the large fraction of the blank sample was similar to that of other environmental samples. A detailed analysis of the data showed that the latter fraction included some of the same ASVs present in the previously processed sample, suggestive of cross-contamination during the DNA extraction process. The possibility of other cross-contaminations between environmental samples was evaluated, but all of the fraction replicates for each sample were of high similarity, and the different fractions for each water sample generally clustered together. We thus concluded that the cross-contamination was limited to the blank sample.

The four samples corresponding to the mock community, one for the small and one for the large fraction, and two for sediments, showed high similarity (Suppl. Fig. 2), in agreement with the fact that all of them contained exactly the same dinoflagellate and parasite community. All reads obtained in the mock community samples, which included several target organisms from this study, belonged to those species added from cultures. Therefore, our metabarcoding method allowed the recovery, amplification and sequencing of target Dinophyceae and Perkinsea. A possible relationship between the number of reads obtained for each organism and the number of cells $mL^{-1}$ added at each sample was evaluated. Firstly, all values were normalized based on the contribution of each species to the total. Next, the relationship between the two variables was determined using a linear regression for each sample (small fraction, large fraction, sediment 1 and sediment 2). No relationship was found ($p > 0.5$) when the values for all eight organisms were processed together, i.e., 4 cultured dinoflagellates species and the hosts present in the parasitoid cultures and the 4 cultured parasitoids (data not shown). The two groups of organisms were then processed independently and the contribution of each species with respect to the total of its group was calculated. For the group of dinoflagellates (Suppl. Fig. 3), the only correlations were found between the number of reads and the cells $mL^{-1}$ of each species in the sediment 2 sample ($p < 0.05$ and $R^2 = 0.94$; Suppl. Fig. 2D). But even in this case, the relation was dominated by the presence of *A. minutum*, which had a strong contribution to cell abundances and to the number of reads. By contrast, the correlations between sequencing reads and cellular abundances of parasitoids (Suppl. Fig. 4) were significant for all samples ($p < 0.05$). The values were lower in the water samples, ($R^2 = 0.84$ in the small fraction and $R^2 = 0.81$ in the large fraction; Suppl. Fig. 4A, B) than in the sediment samples ($R^2 = 0.99$ in sediment 1 and $R^2 = 0.98$ in sediment 2; Suppl. Fig. 4C, D). Even though metabarcoding is considered as a semi-quantitative method, the results obtained for parasitoid suggest that a good relationship between the number of reads and their cellular abundances can be established in results obtained for environmental samples.

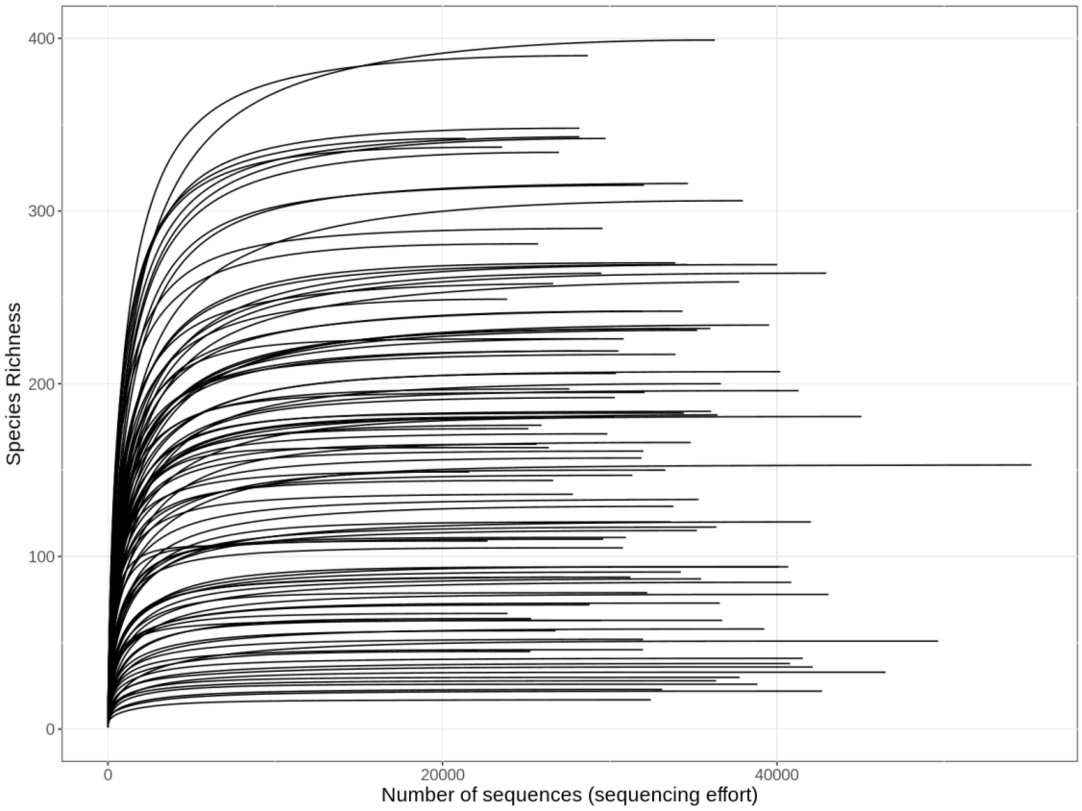


Supplementary Figure 1: Rarefaction curves of all the samples, showing the richness saturation. The horizontal axis indicates the number of sequences (sequencing effort) and the vertical axis the species richness (number of ASVs).

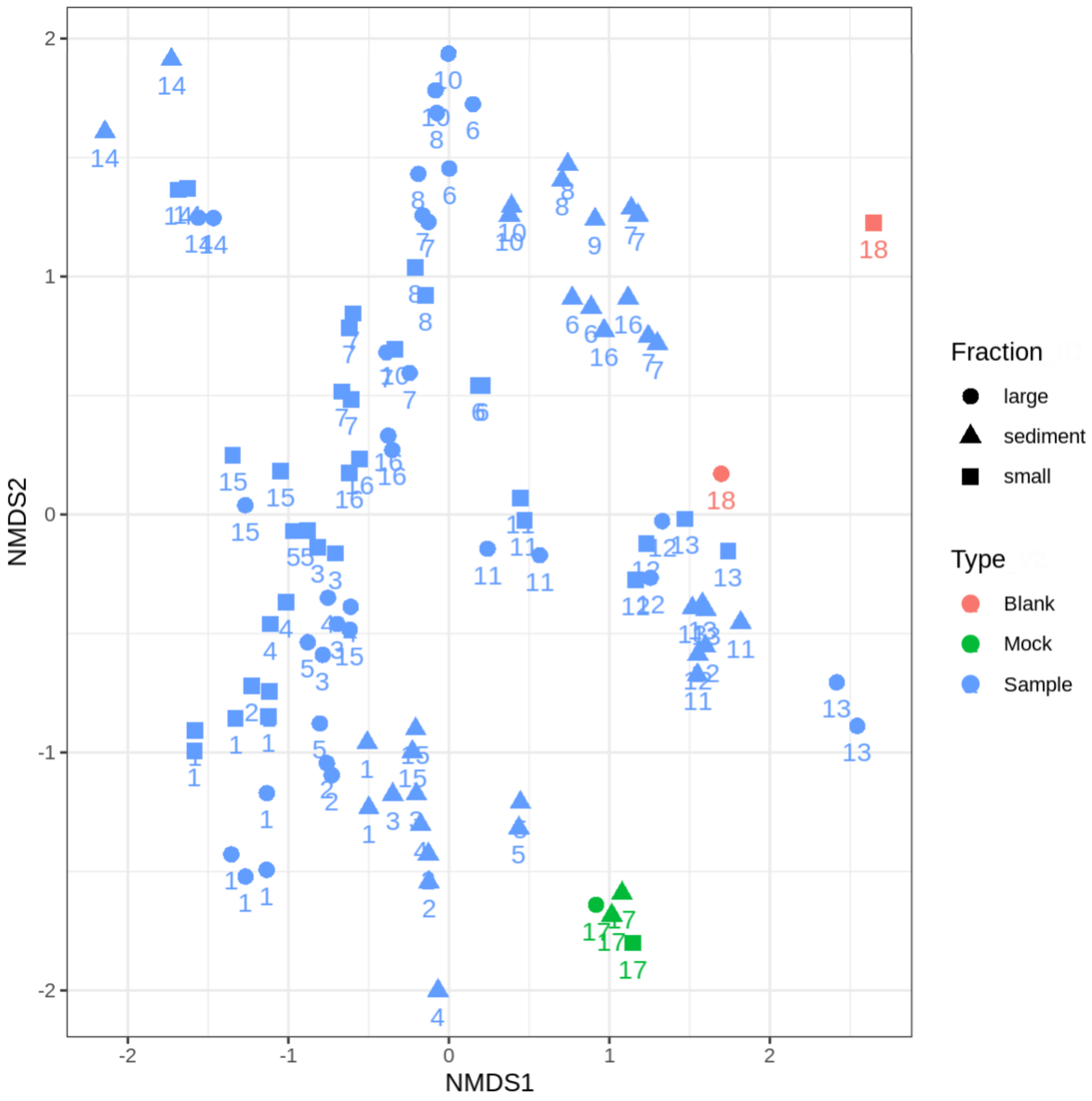


Supplementary Figure 2. Non-metric multidimensional scaling (NMDS) ordination plot, based on Bray-Curtis distances between all the samples including the mocks and blanks. The type of sample is indicated by the different colours (blanks in red, mock in green and samples in blue) and the fraction by the different shapes (small in squares, large in circles and sediments in triangles). The numbers represent the sampling, as described in Table 1.

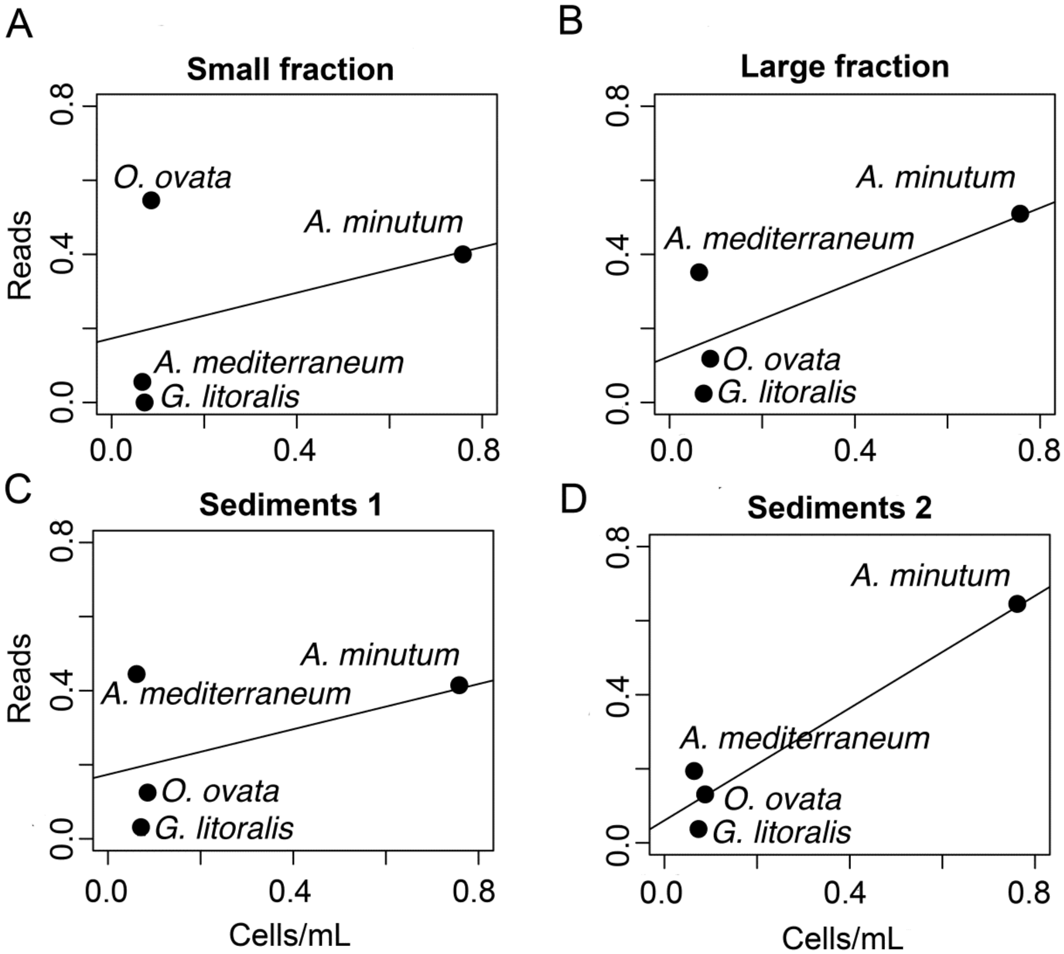


Supplementary Figure 3: Scatterplot of the relation between the normalized contribution of reads (y axis) and cells $mL^{-1}$ (x axis) of each dinoflagellate culture to the total. A) Small fraction (0.8–10 µm). B) Large fraction (10–200 µm). C) Sediment sample 1. D) Sediment sample 2.

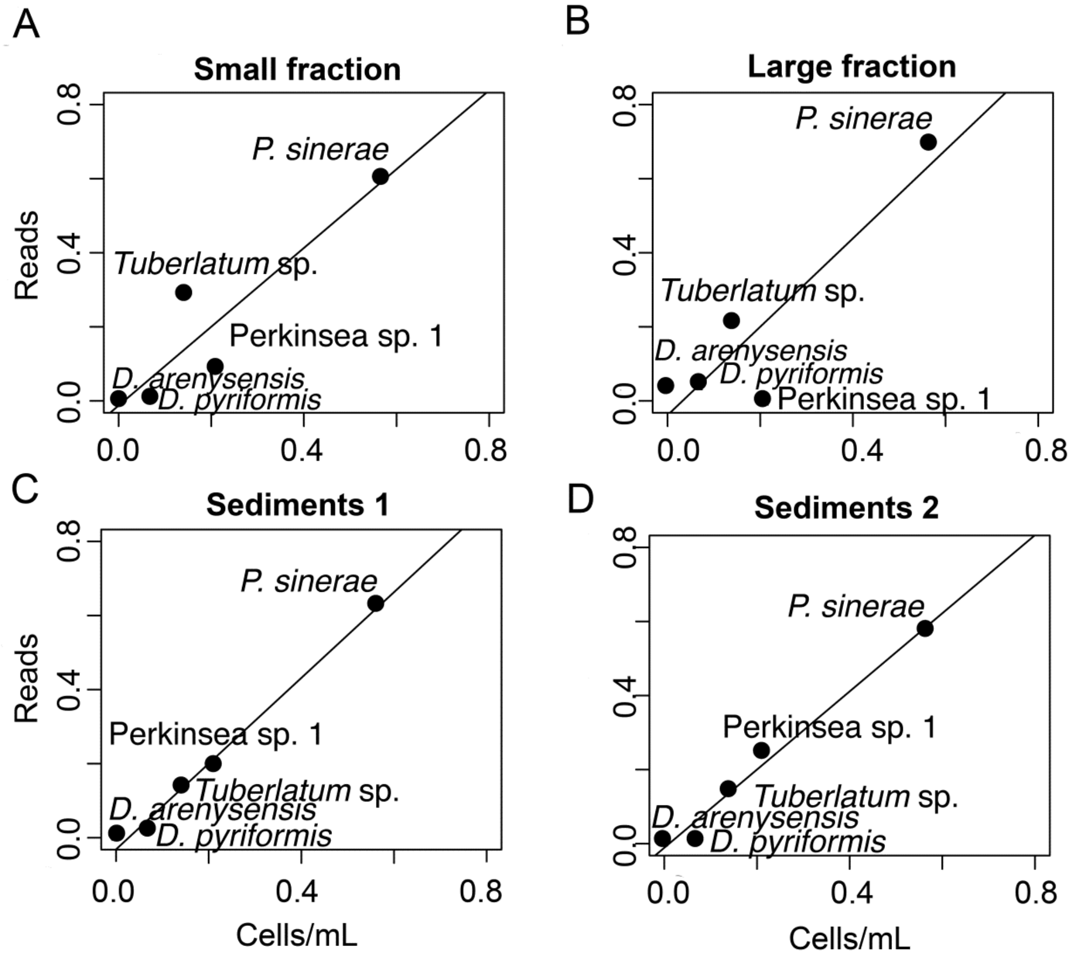


Supplementary Figure 4: Scatterplot of the relation between the normalized contribution of reads (y axis) and cells $mL^{-1}$ (x axis) of each parasitoid culture to the total. A) Small fraction (0.8–10 µm). B) Large fraction (10–200 µm). C) Sediment sample 1. D) Sediment sample 2.

Supplementary Table 1: Relation of metabarcoding samples, including the total number of eukaryotic reads and ASVs obtained, and those corresponding to Dinophyceae and Perkinsea, including the community percentage (%) they represent.

| Sample | | | | Eukaryotes | | Dinophyceae | | | Perkinsea | | |
|---|---|---|---|---|---|---|---|---|---|---|---|
| Code | Location | Replicate | Fraction | Reads | ASVs | Reads | % | ASVs | Reads | % | ASVs |
| CP001 | El Masnou | R1 | small | 27801 | 136 | 9288 | 33.4 | 13 | 0 | 0 | 0 |
| CP002 | El Masnou | R2 | small | 31364 | 147 | 9920 | 31.6 | 13 | 0 | 0 | 0 |
| CP003 | El Masnou | R1 | small | 36732 | 63 | 32712 | 89.1 | 3 | 0 | 0 | 0 |
| CP004 | El Masnou | R2 | small | 40850 | 85 | 36066 | 88.3 | 4 | 0 | 0 | 0 |
| CP005 | Vilanova | R1 | small | 43078 | 78 | 3337 | 7.7 | 12 | 0 | 0 | 0 |
| CP006 | Vilanova | R2 | small | 49637 | 51 | 5162 | 10.4 | 13 | 0 | 0 | 0 |
| CP007 | Llavaneres | R1 | small | 30241 | 181 | 12120 | 40.1 | 52 | 0 | 0 | 0 |
| CP008 | Llavaneres | R2 | small | 33917 | 270 | 16632 | 49 | 49 | 0 | 0 | 0 |
| CP009 | Sant Pol | R1 | small | 33812 | 129 | 17270 | 51.1 | 23 | 95 | 0.28 | 3 |
| CP010 | Sant Pol | R2 | small | 33632 | 120 | 19240 | 57.2 | 24 | 75 | 0.22 | 3 |
| CP011 | Sitges | R1 | small | 27590 | 197 | 2485 | 9 | 14 | 50 | 0.18 | 1 |
| CP012 | Sitges | R2 | small | 30528 | 219 | 2072 | 6.8 | 15 | 0 | 0 | 0 |
| CP013 | Barcelona | R1 | small | 25724 | 281 | 5714 | 22.2 | 36 | 29 | 0.11 | 1 |
| CP014 | Barcelona | R2 | small | 29574 | 290 | 7017 | 23.7 | 36 | 19 | 0.06 | 1 |
| CP015 | Cambrils | R1 | small | 22696 | 109 | 9936 | 43.8 | 21 | 0 | 0 | 0 |
| CP016 | Cambrils | R2 | small | 36444 | 182 | 15140 | 41.5 | 32 | 0 | 0 | 0 |
| CP017 | Muga | R1 | small | 26357 | 163 | 13334 | 50.6 | 25 | 0 | 0 | 0 |
| CP018 | Muga | R2 | small | 25141 | 174 | 13292 | 52.9 | 24 | 0 | 0 | 0 |
| CP019 | Estartit | R1 | small | 29859 | 171 | 7620 | 25.5 | 20 | 0 | 0 | 0 |
| CP020 | Estartit | R2 | small | 26623 | 144 | 5680 | 21.3 | 21 | 0 | 0 | 0 |
| CP021 | Fra Ramon | R1 | small | 25265 | 45 | 4740 | 18.8 | 5 | 0 | 0 | 0 |
| CP022 | Fra Ramon | R2 | small | 40785 | 38 | 12325 | 30.2 | 10 | 0 | 0 | 0 |
| CP023 | Aiguafreda | R1 | small | 23891 | 67 | 18296 | 76.6 | 16 | 0 | 0 | 0 |
| CP024 | Aiguafreda | R2 | small | 35465 | 87 | 27256 | 76.9 | 23 | 0 | 0 | 0 |
| CP025 | L'Arenal | R1 | small | 32082 | 195 | 15394 | 48 | 23 | 0 | 0 | 0 |
| CP026 | L'Arenal | R2 | small | 34434 | 183 | 19228 | 55.8 | 25 | 0 | 0 | 0 |
| CP027 | L'Ametlla | R1 | small | 36065 | 184 | 12982 | 36 | 32 | 0 | 0 | 0 |
| CP028 | L'Ametlla | R2 | small | 31907 | 157 | 14543 | 45.6 | 31 | 2 | 0.01 | 1 |
| CP029 | Pals | R1 | small | 25295 | 64 | 23693 | 93.7 | 18 | 8 | 0.03 | 1 |
| CP030 | Pals | R2 | small | | | | | | | | |
| CP031 | L'Arenal | R1 | small | 29284 | 226 | 6584 | 22.5 | 26 | 47 | 0.16 | 3 |
| CP032 | L'Arenal | R2 | small | 34614 | 269 | 12021 | 34.7 | 32 | 15 | 0.04 | 2 |
| CP033 | Cubelles | R1 | small | 34249 | 91 | 30233 | 88.3 | 26 | 0 | 0 | 0 |
| CP034 | Cubelles | R2 | small | 42037 | 120 | 36448 | 86.7 | 34 | 0 | 0 | 0 |

| | | | | | | | | | | | |
|---|---|---|---|---|---|---|---|---|---|---|---|
| CP037 | El Masnou | R1 | large | 39238 | 58 | 37099 | 94.5 | 13 | 0 | 0 | 0 |
| CP038 | El Masnou | R2 | large | 46493 | 33 | 45974 | 98.9 | 12 | 0 | 0 | 0 |
| CP039 | El Masnou | R1 | large | 36367 | 28 | 36118 | 99.3 | 12 | 0 | 0 | 0 |
| CP040 | El Masnou | R2 | large | 42696 | 22 | 42554 | 99.7 | 8 | 0 | 0 | 0 |
| CP041 | Vilanova | R1 | large | 29488 | 63 | 13071 | 44.3 | 26 | 0 | 0 | 0 |
| CP042 | Vilanova | R2 | large | 36585 | 73 | 12200 | 33.3 | 29 | 0 | 0 | 0 |
| CP043 | Llavaneres | R1 | large | 55217 | 153 | 46776 | 84.7 | 60 | 27 | 0.05 | 2 |
| CP044 | Llavaneres | R2 | large | 32018 | 161 | 24905 | 77.8 | 63 | 49 | 0.15 | 1 |
| CP045 | Sant Pol | R1 | large | 31988 | 46 | 27716 | 86.6 | 28 | 250 | 0.78 | 2 |
| CP046 | Sant Pol | R2 | large | 31998 | 52 | 27257 | 85.2 | 29 | 169 | 0.53 | 3 |
| CP047 | Sitges | R1 | large | 40195 | 207 | 3486 | 8.7 | 25 | 66 | 0.16 | 2 |
| CP048 | Sitges | R2 | large | 45045 | 181 | 11566 | 25.7 | 31 | 60 | 0.13 | 3 |
| CP049 | Barcelona | R1 | large | 30307 | 192 | 16743 | 55.2 | 30 | 14 | 0.05 | 1 |
| CP050 | Barcelona | R2 | large | 42946 | 264 | 23921 | 55.7 | 32 | 33 | 0.08 | 1 |
| CP051 | Cambrils | R1 | large | 40674 | 94 | 31948 | 78.5 | 26 | 0 | 0 | 0 |
| CP052 | Cambrils | R2 | large | 40109 | 94 | 35492 | 88.5 | 34 | 0 | 0 | 0 |
| CP053 | Muga | R1 | large | 27287 | 88 | 22292 | 81.7 | 23 | 0 | 0 | 0 |
| CP054 | Muga | R2 | large | 29606 | 110 | 22300 | 75.3 | 28 | 0 | 0 | 0 |
| CP055 | Estartit | R1 | large | 33333 | 150 | 25567 | 76.7 | 35 | 4 | 0.01 | 1 |
| CP056 | Estartit | R2 | large | 30974 | 111 | 24106 | 77.8 | 37 | 0 | 0 | 0 |
| CP057 | Fra Ramon | R1 | large | 28794 | 72 | 14921 | 51.8 | 12 | 0 | 0 | 0 |
| CP058 | Fra Ramon | R2 | large | 32247 | 79 | 20157 | 62.5 | 16 | 0 | 0 | 0 |
| CP059 | Aiguafreda | R1 | large | 37774 | 30 | 37401 | 99 | 18 | 0 | 0 | 0 |
| CP060 | Aiguafreda | R2 | large | 41546 | 41 | 39668 | 95.5 | 23 | 0 | 0 | 0 |
| CP061 | L’Arenal | R1 | large | 30791 | 105 | 11080 | 36 | 38 | 0 | 0 | 0 |
| CP062 | L’Arenal | R2 | large | 35312 | 133 | 16922 | 47.9 | 64 | 0 | 0 | 0 |
| CP063 | L’Ametlla | R1 | large | 26766 | 57 | 26096 | 97.5 | 26 | 0 | 0 | 0 |
| CP064 | L’Ametlla | R2 | large | 31234 | 88 | 27447 | 87.9 | 35 | 0 | 0 | 0 |
| CP065 | Pals | R1 | large | 32463 | 17 | 32406 | 99.8 | 10 | 0 | 0 | 0 |
| CP066 | Pals | R2 | large | 33131 | 23 | 33044 | 99.7 | 16 | 0 | 0 | 0 |
| CP067 | L’Arenal | R1 | large | 36384 | 117 | 31888 | 87.6 | 46 | 0 | 0 | 0 |
| CP068 | L’Arenal | R2 | large | 35210 | 115 | 32125 | 91.2 | 45 | 16 | 0.05 | 1 |
| CP069 | Cubelles | R1 | large | 42138 | 36 | 35354 | 83.9 | 16 | 20 | 0.05 | 1 |
| CP070 | Cubelles | R2 | large | 38837 | 26 | 30101 | 77.5 | 11 | 11 | 0.03 | 1 |
| CP073 | El Masnou | R1 | sediment | 34838 | 166 | 23643 | 67.9 | 21 | 0 | 0 | 0 |
| CP074 | El Masnou | R1 | sediment | 41299 | 196 | 18173 | 44 | 38 | 0 | 0 | 0 |
| CP075 | Vilanova | R1 | sediment | 23875 | 249 | 8343 | 34.9 | 49 | 0 | 0 | 0 |
| CP076 | Vilanova | R2 | sediment | 26618 | 258 | 8483 | 31.9 | 52 | 0 | 0 | 0 |
| CP077 | Llavaneres | R1 | sediment | 33613 | 232 | 5669 | 16.9 | 14 | 5 | 0.01 | 1 |
| CP078 | Llavaneres | R2 | sediment | 36014 | 232 | 6858 | 19 | 16 | 0 | 0 | 0 |

| | | | | | | | | | | | |
|---|---|---|---|---|---|---|---|---|---|---|---|
| CP079 | Sant Pol | R1 | sediment | 25924 | 176 | 17268 | 66.6 | 35 | 191 | 0.74 | 4 |
| CP080 | Sant Pol | R2 | sediment | 35220 | 231 | 22472 | 63.8 | 41 | 462 | 1.31 | 4 |
| CP081 | Sitges | R1 | sediment | 34355 | 242 | 3909 | 11.4 | 17 | 0 | 0 | 0 |
| CP082 | Sitges | R2 | sediment | 33938 | 217 | 5975 | 17.6 | 13 | 0 | 0 | 0 |
| CP083 | Barcelona | R1 | sediment | 29763 | 342 | 4683 | 15.7 | 24 | 20 | 0.07 | 2 |
| CP084 | Barcelona | R2 | sediment | 36291 | 399 | 6745 | 18.6 | 27 | 36 | 0.1 | 2 |
| CP085 | Cambrils | R1 | sediment | 37739 | 259 | 13945 | 37 | 29 | 0 | 0 | 0 |
| CP086 | Cambrils | R2 | sediment | 36645 | 200 | 18102 | 49.4 | 25 | 2 | 0.01 | 1 |
| CP087 | Estartit | R1 | sediment | 29503 | 264 | 3401 | 11.5 | 28 | 154 | 0.52 | 1 |
| CP088 | Estartit | R2 | sediment | 6158 | 106 | 91 | 1.5 | 4 | 22 | 0.36 | 1 |
| CP089 | Muga | R1 | sediment | 28190 | 348 | 3697 | 13.1 | 42 | 51 | 0.18 | 5 |
| CP090 | Muga | R2 | sediment | 28692 | 390 | 3530 | 12.3 | 35 | 37 | 0.13 | 3 |
| CP091 | Fra Ramon | R1 | sediment | 25627 | 165 | 346 | 1.4 | 4 | 32 | 0.12 | 1 |
| CP092 | Fra Ramon | R2 | sediment | 21613 | 149 | 134 | 0.6 | 6 | 52 | 0.24 | 1 |
| CP093 | Aiguafreda | R1 | sediment | 34679 | 316 | 10568 | 30.5 | 29 | 0 | 0 | 0 |
| CP094 | Aiguafreda | R2 | sediment | 32049 | 315 | 11362 | 35.5 | 24 | 0 | 0 | 0 |
| CP095 | L’Arenal | R1 | sediment | 28191 | 343 | 2279 | 8.1 | 30 | 3 | 0.01 | 1 |
| CP096 | L’Arenal | R2 | sediment | 26968 | 334 | 2036 | 7.5 | 28 | 8 | 0.03 | 1 |
| CP097 | L’Ametlla | R1 | sediment | 21385 | 342 | 2458 | 11.5 | 16 | 0 | 0 | 0 |
| CP098 | L’Ametlla | R2 | sediment | 23562 | 337 | 2044 | 8.7 | 15 | 0 | 0 | 0 |
| CP099 | Pals | R1 | sediment | 40005 | 269 | 11307 | 28.3 | 23 | 0 | 0 | 0 |
| CP100 | Pals | R2 | sediment | 39537 | 234 | 13226 | 33.5 | 26 | 40 | 0.1 | 2 |
| CP101 | L’Arenal | R1 | sediment | 30373 | 206 | 3981 | 13.1 | 21 | 0 | 0 | 0 |
| CP102 | L’Arenal | R2 | sediment | 26034 | 180 | 2369 | 9.1 | 16 | 0 | 0 | 0 |
| CP103 | Cubelles | R1 | sediment | 31978 | 242 | 3615 | 11.3 | 18 | 9996 | 31.26 | 4 |
| CP104 | Cubelles | R2 | sediment | 28332 | 219 | 3016 | 10.6 | 16 | 8076 | 28.5 | 3 |
| CP105 | Sa Riera | R1 | sediment | 37967 | 306 | 24791 | 65.3 | 35 | 89 | 0.23 | 1 |
| CP106 | Cubelles | R1 | sediment | 30836 | 226 | 3368 | 10.9 | 16 | 8953 | 29.03 | 3 |

Supplementary Table 2: List of Perkinsea parasitoid cultures established, including details of its isolation source, molecular rDNA sequences obtained, and dinoflagellate hosts used for maintaining the cultures.

| Species | Location | Date | 18S rDNA | Host | Strain | Origin | Year of isolation |
|---|---|---|---|---|---|---|---|
| *Parvilucifera sinerae* | Barcelona | 28-mar-19 | MT606014 | *Alexandrium minutum* | Arenys | Arenys (Catalan coast) | 2018 |
| *Parvilucifera sinerae* | Cambrils | 15-may-19 | MT606015 | *Alexandrium minutum* | Arenys | Arenys (Catalan coast) | 2018 |
| *Parvilucifera sinerae* | Pals | 4-jul-19 | MT606016 | *Gymnodinium litoralis* | UNISS1 | Sardinia (Mediterranean Sea) | 2010 |
| *Dinovorax pyriformis* | Llavaneres | 25-jul-18 | MT606012 | *Ostreopsis* sp. | OOPM18 | Llavaneres (Catalan coast) | 2018 |
| *Dinovorax pyriformis* | Sitges | 7-ago-18 | MT606013 | *Ostreopsis* sp. | OOPM18 | Llavaneres (Catalan coast) | 2018 |
| *Dinovorax pyriformis* | Cubelles | 27-ago-19 | MT606011 | *Ostreopsis* sp. | OOPM18 | Llavaneres (Catalan coast) | 2018 |
| *Tuberlatum* sp. | Sant Pol | 25-jul-18 | MT606017 | *Alexandrium taylorii* | VGO703 | Alfacs (Catalan coast) | 2003 |
| | | | | *Alexandrium minutum* | Arenys | Arenys (Catalan coast) | 2018 |
| *Perkinsea* sp. 1 | Sant Pol | 25-jul-18 | MN721815 | *Alexandrium taylorii* | VGO703 | Alfacs (Catalan coast) | 2003 |
| | | | | *Alexandrium minutum* | Arenys | Arenys (Catalan coast) | 2018 |
